# Protected areas enhance avian food webs

**DOI:** 10.1101/2023.11.30.569404

**Authors:** Lucie Thompson, Núria Galiana, Konstans Wells, Miguel Lurgi

**Affiliations:** Department of Biosciences, Swansea University, Singleton Park, SA2 8PP, UK; Department of Biogeography and Global Change, National Museum of Natural Sciences (CSIC), 28006 Madrid, Spain

## Abstract

Restoring and conserving habitat and the species they shelter has become a primary focus to mitigate the current extinction crisis. Setting aside land designated as protected areas (PAs) is an efficient way of achieving these aims. This strategy has been proven to enhance different aspects of species richness and abundance across ecosystems^1–4^. However, to truly understand the effects of global environmental change on biodiversity, and the efficiency of our mitigation measures, we must account for one of its fundamental dimensions: species interactions. Here we show that PAs enhance avian food webs across Europe by protecting key network and species traits. Using 376,556 observational records of 509 bird species from citizen science databases distributed across 45 networks of PAs, we found beneficial effects of protection on 10 out of 13 food web properties on an average of 25.9% of sites. PAs enhance food webs by harbouring large top predators, in turn increasing the length of biomass flow paths from basal to top species. Furthermore, we link these beneficial effects to environmental drivers and PA designations. PA benefits were augmented by specific protection goals such as European Directives for conservation. This study provides evidence for the effectiveness of PAs as a strategy to preserve fundamental aspects of biodiversity beyond species richness. We anticipate our study to be a starting point for the development of comprehensive frameworks to assess the critical role of PAs in safeguarding biodiversity worldwide. Improving the mapping of species occurrences and ecological interactions across the globe will is fundamental to develop optimal strategies for establishing networks of PAs aimed at protecting all aspects of ecosystem diversity.

## Introduction

Around twelve thousand years ago, with the expansion of human populations, the Earth started to transition into a period that we now know as the Anthropocene. Due to anthropogenic change, our world has, since then, experienced hundreds of species extinctions and range contractions. This has resulted in a global collapse of biodiversity and species abundance, with fundamental repercussions for the insidious extinctions of ecological interactions that glue communities together^5,6^. The selective loss of species with particular traits such as large species^7–10^, mammals^11^, and top predators ^12^, has caused a fundamental rearrangement of natural communities and food webs.

Protected areas (PAs), by setting aside natural areas and shielding them from some or all human pressure^13,14^, represent the main conservation strategy to slowing down the biodiversity crisis. They have been proven effective at slowing down species, functional and phylogenetic diversity loss, as well as population declines^1–4,15,16^. Nonetheless, if we are to fully understand the effects of anthropogenic changes on biodiversity, and therefore the effectiveness of PAs at ameliorating them, we need to consider their influence on biotic interactions. Biotic interactions should be systematically included in biodiversity assessments as they are a key component of biological diversity.

Ecological communities are more than the sum of their parts. They are complex systems of interacting species that create emergent patterns and functions^8,17^. The consideration of networks of ecological interactions, such as food webs, when investigating natural ecosystems, is fundamental to our understanding of community stability and ecosystem function. The structure of biotic interactions can inform on a community’s ability to withstand anthropogenic perturbations such as species loss or biomass disturbance, and their cascading effects^18,19^, and to provide crucial ecosystem services^20^. As such, our ability to understand the structure of ecological networks, and predict the dynamics emerging from it, is fundamental to unveil community assembly processes, playing thus a crucial role in devising conservation strategies, including the establishment of PAs ^17,21^.

Gradients of anthropogenic disturbance negatively impact the shape of food webs. Species richness, the average length of food chains, and the fraction of omnivorous species decrease along gradients of habitat degradation^22–24^. Concomitant consequences of these changes on food webs include a reduction in their compartmentalisation, due to a relative increase in trophic interactions when species are lost^23,25,26^, but see^18,27^. Food webs also become shorter and less pyramidal, shifting towards a ‘flatter’ structure with more, but smaller, top predator species due to the loss of the more vulnerable, larger species^7–10,27,28^. Although species-focused assessments of PA efficacy have provided useful insights, it remains unknown to what extent are PAs able to enhance species interactions networks. Here we ask three fundamental questions: first, whether PAs enhance food web structure compared with their immediate vicinity; second, whether the influence of PAs is different across food web features; and third, what are the drivers of these differences.

We assessed the ability of PAs to mitigate the impacts of ecosystem degradation on food webs across Europe by combining the two globally largest citizen science databases of species occurrences, GBIF^29^ and eBird^30^, with a comprehensive, taxonomically-resolved food web of tetrapod species across Europe^31^. We applied a very restrictive filtering on this occurrence data to remove representation biases across regions and species, resulting in 369,799 individual records of 509 bird species across 45 networks of PAs and surrounding areas in Europe. We built local avian food webs at the 10×10km cell level and quantified 15 features characterising food web structure and their species. By contrasting these features between PAs and unprotected surrounding areas using a bootstrapped modelling approach to ensure homogeneity of sampling, we examined the effects of protection on biotic interactions. Using a multivariate modelling framework, we explored how this contrast (i.e. protection outcome) varied across types of protected areas and their environmental context.

### Mixed effects of PAs on food webs

Using 233,771 survey events reporting 376,556 observational records of 509 bird species across 45 PA networks composed of an average of 7 (sd = 6, min = 1, max = 123) PAs and their surrounding areas, altogether called sites hereafter (Fig. 1; Supplementary Data 1 for list of sites and their characteristics), we built local food webs at the 10×10 km grid cell-level for each map cell within PAs and surrounding areas. We chose cells randomly for comparison according to the smallest number of cells between the PA and the outside to avoid sampling bias, yielding an average of 60 grid cells per site (sd = 33, min = 30, max = 198). We repeated this process 300 times to ensure accurate and unbiased quantification of the contrasts.

**Fig. 1.**
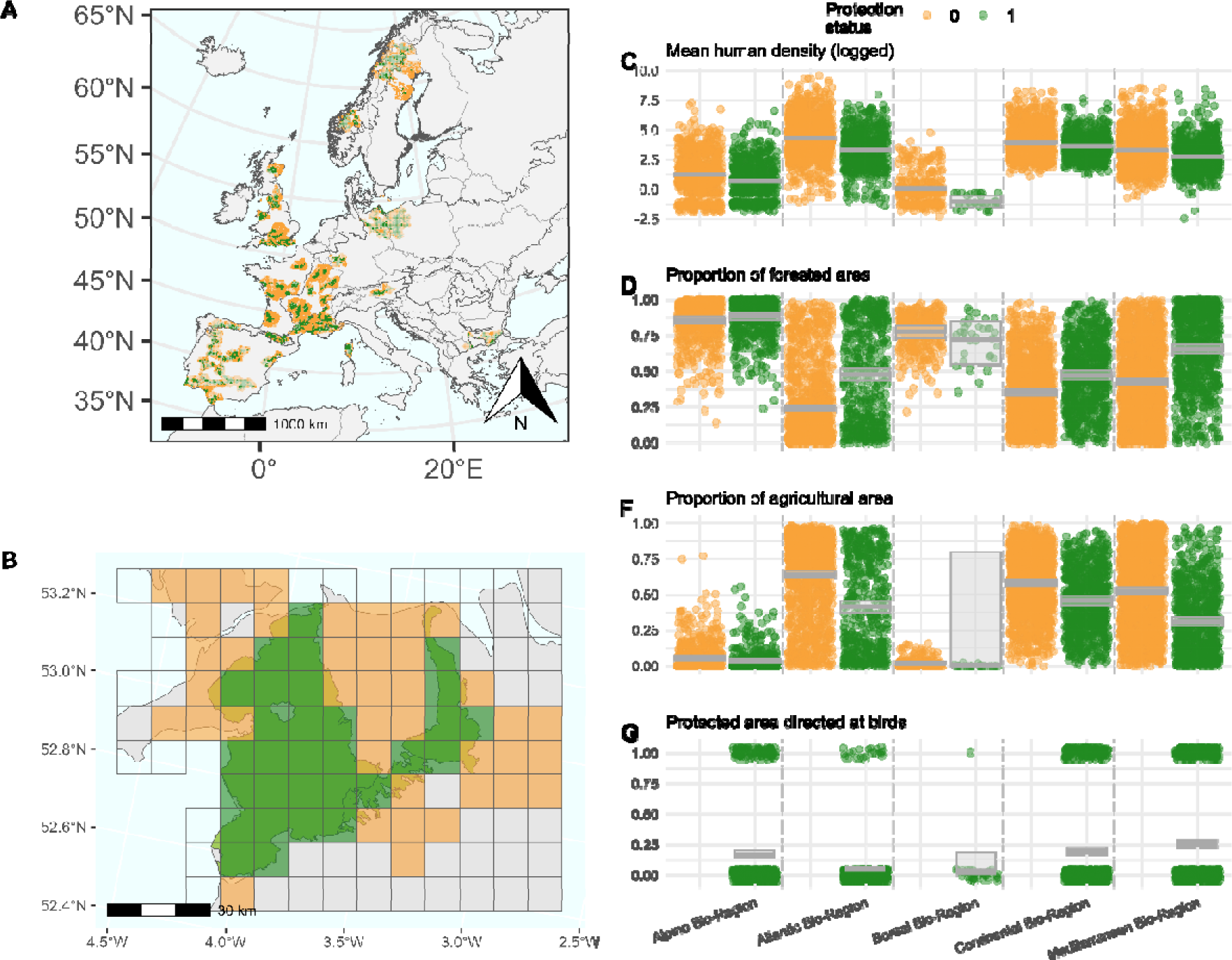
Location of PAs (green) and unprotected surrounding areas (orange) studied across Europe, and corresponding levels of anthropogenic disturbance. The map in (A) shows the 45 PA networks studied. Each site (continuous areas of green and orange grid cells) represents a grouping of connected PAs (less than 1 km apart) within the same biogeographical region and established before 2015. Transparency of the grid cells distinguishes cells used for analysis (subset dataset – 4591 grid cells; darker – Supplementary Data 2) - and those only used to describe the environmental characteristics of the sites (whole dataset – 8932 grid cells; lighter – Supplementary Data 3). Lighter grid cells were left out of the biodiversity analyses due to low sampling effort. Each site comprises protected (green) and unprotected (orange) 10×10km grid cells. Unprotected grid cells lay within 100km of the protected ones. Grey boundaries represent country boundaries and black boundaries represent biogeographical bioregion boundaries (<www.eea.europa.eu>). Protected areas (PAs) were extracted from the World Database on Protected Areas (WDPA)^34^. (B) shows a sample location in Wales, United Kingdom, illustrating the local scale detail of resolution of map grid cells considered within biodiversity and environmental records. Only coloured grid cells were used for analysis (sufficient survey effort), but all grid cells (coloured and transparent) were used to evaluate environmental characteristics of the site. (C-F) Show the difference on Anthropogenic impact in protected vs unprotected land across the 5 bioregions represented in our study. The fraction of the PAs especially designated for the protection of bird species (G) is related to the efficacy of PAs at protecting avian food webs (Fig. 4). Horizontal bars and shaded area on the data distributions represent the estimate and 95% confidence intervals of the linear (C) or logistic regressions (D-F) of the environmental characteristic as a function of the interaction between bioregion and protection status.

We found that 63% of the sites did not show a significant contrast in food web metrics between protected and non-protected communities (Fig. 2). Nonetheless, a substantial number of sites did show a higher species richness (33% of sites), longer mean food chain length (MFCL) (29% of sites), number of species at basal and intermediate positions of the food webs (31% and 38% of sites respectively) and mean diet breadth (24% of sites) (Fig. 2). In empirical and theoretical food webs, longer MFCL and more intermediate species are typically associated with higher productivity (e.g. productive habitats)^32^ or larger and better connected sites^33^, i.e. habitats that can host more species, but see^25^. Intermediate species act as both resources and consumers, linking top and basal species, and thus creating indirect interactions and facilitating the flow of energy from lower to higher trophic levels^8^.

**Fig. 2.**
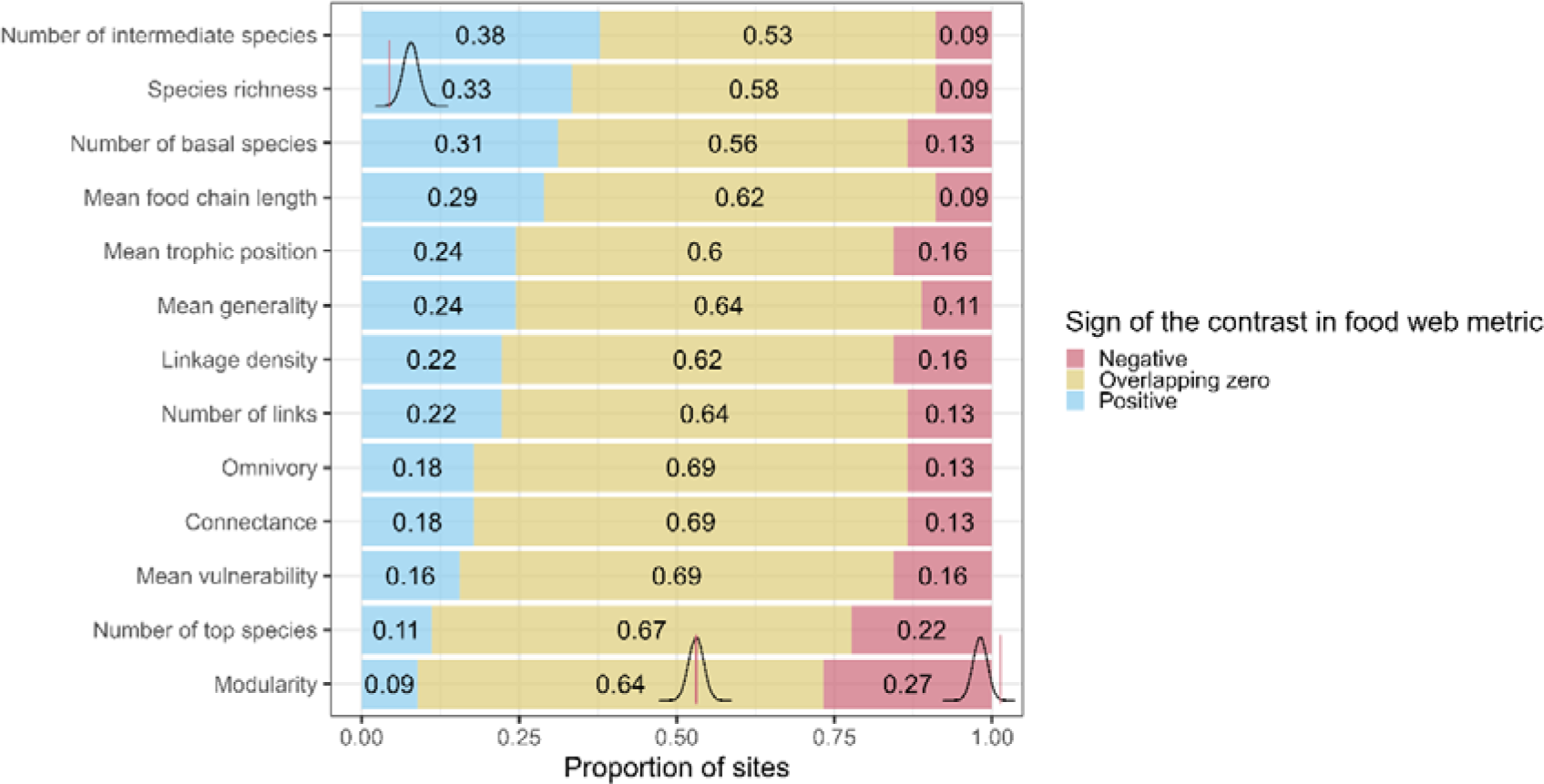
Differential effects of PA protection across food web properties. Proportion of sites with a positive (blue), negative (red), or neutral (yellow) effect of PAs (vs neighbouring unprotected areas) in a diverse array of 13 food web metrics. The numbers display the proportion of sites among the 45 sites studied here (Fig. 1) showing the corresponding sign of the effect on each given food web property. The distribution diagrams (black bell-shaped curves overlaid on plots) illustrate the three hypothetical scenarios (positive, negative, neutral distributions in contrasts in metrics) with respect to zero (red vertical line). Contrasts / effects between PAs vs unprotected areas were quantified as the average contrast in each property between pairs of randomly sampled grid cells inside vs outside of each PA (average 60 pairs of grid cells per site +/- 33 pairs). The random selection of pairs of grid cells was bootstrapped 300 times, thus generating a distribution of average contrasts across grid cells inside vs outside per site (see Methods). The fraction of sites within each contrast level was quantified as the proportion of sites for which the 95% quantiles of the distribution of contrasts overlapped zero (neutral, yellow), were larger (positive, blue), or smaller (negative, red) than 0.

### Top species connect food chains inside PAs

Compartmentalisation and number of top species were found to be smaller within PAs across 27% (vs 9% larger) and 22% (vs 11% larger) of sites, respectively (Fig. 2). PAs holding significantly less top species also displayed longer MFCLs and higher mean trophic position than the surrounding unprotected areas (Pearson r = -0.77 for MFCL vs number of top species). This suggests that higher, and possibly larger, top predators link many food chains together by predating on smaller predators that occupy top positions in their respective chains when higher top predators are absent. This in turns reduces compartmentalisation by linking different parts of the ecosystem (i.e. different food chains).

To further explore this hypothesis, and the role of top species in protected vs unprotected areas, and thus find a mechanistic link between a loss of compartmentalisation and the number of top species within PAs, we measured the contrast in body mass of top species across sites. A high proportion of sites did display higher mean body mass of intermediate and top species inside PAs (36% and 27% of sites respectively vs only 7% and 11% of sites displaying the opposite trend, respectively; Fig. 3). This finding shows the ability of PAs of harbouring larger species that are important for different ecosystem functions.

**Fig. 3.**
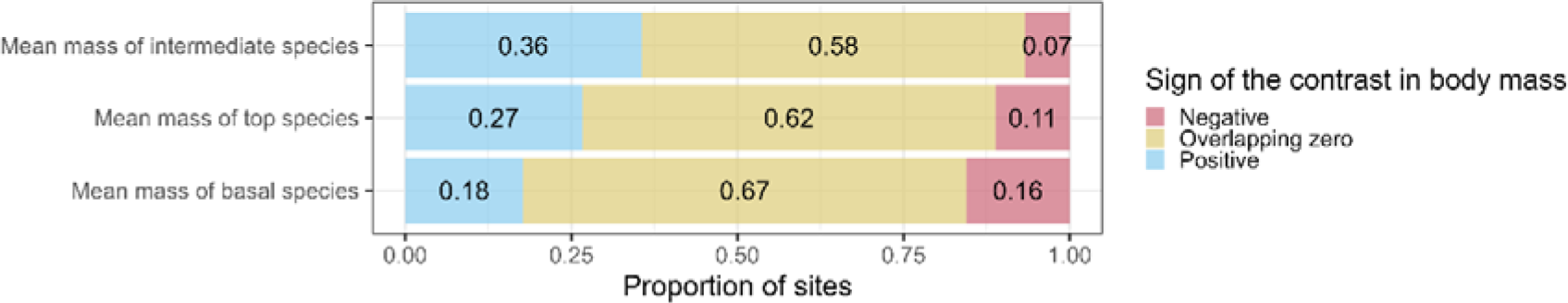
Differential effects of PA protection on species body size across trophic levels. Proportion of sites with a positive (blue), negative (red), or neutral (yellow) effects of PAs (vs neighbouring unprotected areas) on the average body size of adult individuals from the species comprising the food web classified into 3 different trophic levels according to the position they occupy within the food web: top, intermediate or basal. The numbers display the proportion of sites among the 45 sites studied here (Fig. 1) displaying the corresponding sign and significance of the effect / contrast on body size of species within the corresponding trophic level. The fraction of sites within each contrast level was quantified as the proportion of sites for which the 95% quantiles of the distribution of contrasts overlapped zero (neutral, yellow), were larger (positive, blue), or smaller (negative, red) than 0.

Together these findings indicate that PAs enhance food webs by harbouring larger top predators capable of linking ecosystem compartments and making food webs longer and more pyramidal, thus enhancing the functioning and stability of ecosystems ^35^.

### Environmental context drives PA effects

To unveil the drivers behind the large variability in effects on food web metrics across sites, we investigated an array of potential drivers describing environmental and geographical features, as well as purpose of protection across sites. We assessed the relationships between the mean contrast in food web properties between the inside and the outside of PAs for each site to characteristics of PAs and their unprotected surroundings, and their difference. Stepwise variable selection based on Bayes Information Criterion (BIC) revealed significant drivers of change for 9 of the 13 food web metrics and mean body size of top species. The effect of PAs on MFCL, mean body size and number of top species, and mean generality varied consistently with our set of drivers (respective adjusted R² = 0.55, 0.27, 0.26, and 0.22; Fig. 4). In contrast, the number of intermediate and basal species, number of links, linkage density, connectance and mean generality showed less consistent relationships (adjusted R² between 0.14 and 0.09; Fig. 4).

**Fig. 4.**
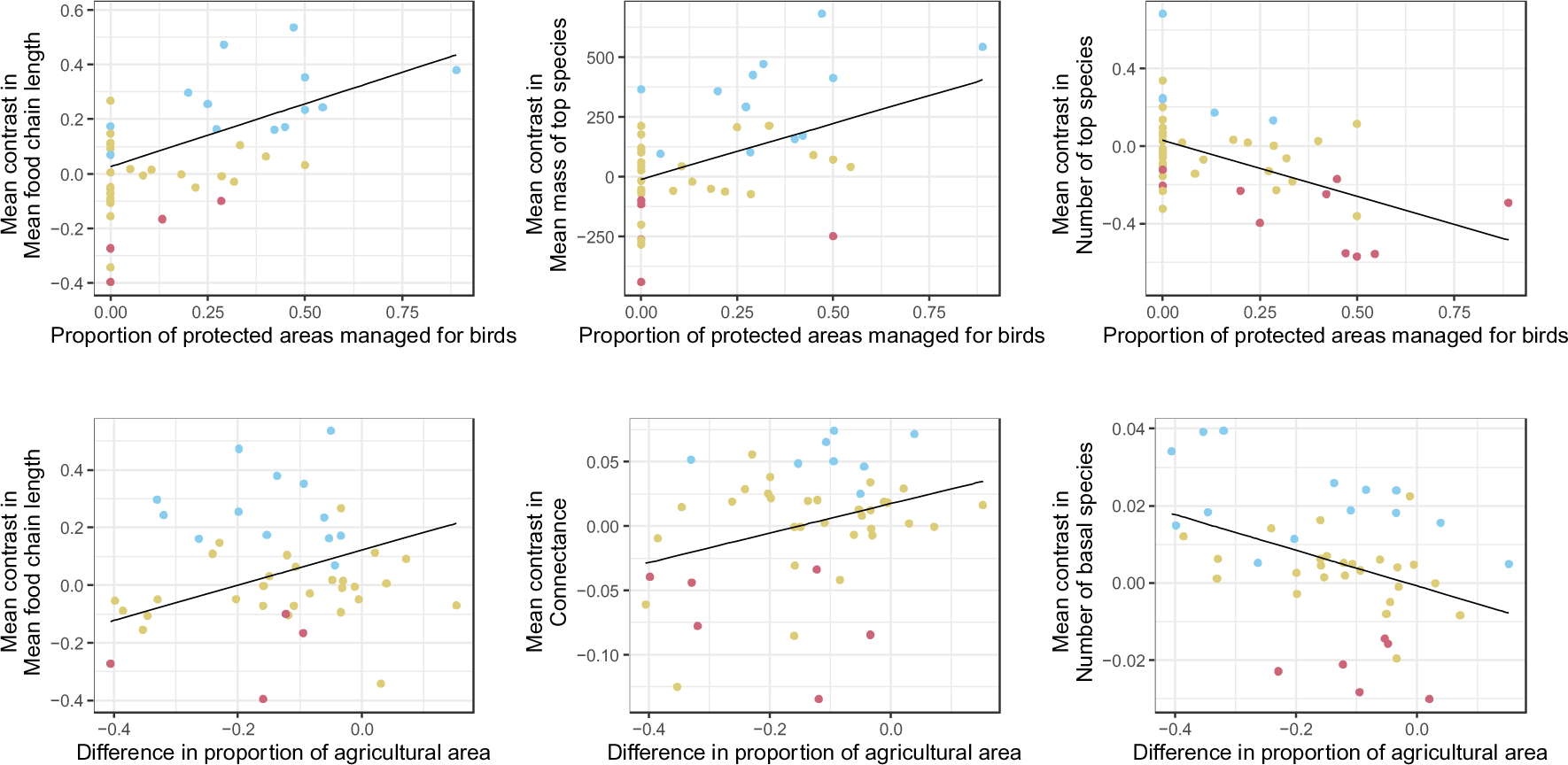
Abiotic conditions drive the effects of PAs across food web properties. We assessed the partial effects of proportion of PAs managed for birds and the difference in proportion of agricultural area inside vs outside of the protected area (among other environmental drivers -see Methods and Supplementary Figure 1 for full set of environmental predictors-) on the mean contrast in food web properties and species body size. Points on scatter plots represent the average contrast across 300 bootstrapped iterations of randomly chosen cells inside vs outside of each of the PAs analysed here (n = 45 sites, average 60 grid map cells per site). See Fig. 1 and Methods for details on data and bootstrapping methodology. Lines and shaded area around them show the partial linear regression with 95% confidence interval (see Supplementary Figure 1 for all scatter plots, and Supplementary Table 1 for full set of coefficients and significance for the scaled linear models). Each linear regression also included one extra variable to statistically account for the variable proportion of PAs across sites (see Supplementary Tables 1 and 2 for the full scaled and non-scaled results, respectively). Point colours indicate the sign of the contrast in food web metric: significantly positive contrasts are highlighted in blue (i.e. larger food web property inside than outside protected areas – see Fig. 2), while negative contrasts are shown red (i.e. larger food web property outside than inside protected areas). Neutral contrast (i.e. no effect) are shown in yellow.

PAs specifically managed for birds (Natura 2000 bird directive sites and Ramsar sites for wetlands) host food webs with longer MFCL, a smaller number of top species but with larger body size and more intermediate species (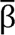 = 0.099, se = 0.022, p < 0.001, df = 37; 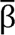 = -0.124, se = 0.031, p < 0.001, df = 42; 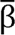 = 100.594, se = 30.709, p < 0.01, df = 41; and 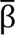 = 0.053, se = 0.018, p < 0.01, df = 42, respectively – see Supplementary Table 1 for full set of coefficients), revealing that the relatively small positive effects of PAs on avian food webs (Fig. 2) are mainly driven by the purpose for which PAs are designated. Natura 2000 areas of protection have been created with the aim to fulfil European Directive 2009/147/EC of the 30^th^ of November 2009 for the conservation of wild bird populations^36^, to preserve, maintain or restore a sufficient diversity and area of habitats for the conservation of all species of birds, with special conservation measures for the habitats of certain species to ensure their survival and reproduction in their area of distribution. The Ramsar protocol aims to promote the conservation of wetlands and waterfowls^37^. Our results suggest that both Natura 2000 and Ramsar designations are fundamental not only to protect bird diversity but also to the conservation of biotic interactions and the structure of the ecological networks they shape.

The difference in proportion of agricultural areas between PAs and their unprotected surroundings also related positively with the contrast in MFCL (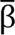 = 0.082 se = 0.03, p< 0.01, df = 37), connectance (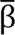 = 0.015, se = 0.007, p< 0.05), linkage density (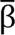 = 0.07, se = 0.04, p< 0.05, df = 42) and mean vulnerability (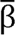 = 0.07, se = 0.037, p< 0.1, df = 42). Conversely, the contrast in number of basal species was favoured in sites with less agricultural area inside than outside (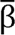 = - 0.006, se = 0.002, p-value < 0.01, df = 42). Although the effect sizes across different models are not comparable, the effect size of the difference in agricultural area on the contrast in MFCL was the smallest among all the variables selected.

### The influence of external disturbance

Conditions outside protected areas influence the outcome of protection. The difference in human density (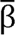 = -0.13, se = 0.054, p < 0.05, df = 37) and the proportion of urban habitat outside PAs (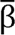 = -0.246, se = 0.079, p < 0.01, df = 37) both correlated negatively with the contrast in MFCL. Thus, when PAs surroundings are subject to higher anthropogenic disturbances, we observe a positive effect of the PA, where pressure is reduced. This was also the case for the relation between higher human density outside PAs and the average number of feeding interactions across species (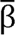 = -0.29, se = 0.119, p value < 0.05, df = 42), which tended to homogenise between the inside and outside when human pressure was high (Supplementary Figure 1). The lack of refugia outside of PAs may restrict habitats available for protected species and thus constraining resource availability for the species, and ultimately the number of species that can be supported across the whole site.

Intrinsic features of PAs are also relevant to the maintenance of complex food webs. We found that the contrast in MFCL was correlated with the remoteness of the site (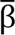 = -0.17, se = 0.045, p-value < 0.001, df = 37), with the outside unprotected vicinity of PAs in more remote sites, displaying longer food chains than their protected parts. This result is surprising, as we expected remote sites to be completely homogeneous, but it suggests that the establishment of PAs in remote areas, where they are commonly set due to human convenience ^38^, might not be a conservation priority. Finally, we observed that PAs hosting larger top predators occurred in sites with higher diversity of land cover types (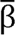 = 66.93, se = 34.61, p-value < 0.01, df = 42). This highlights the importance of maintaining habitat heterogeneity for ecosystem health.

### Enhancing biodiversity beyond species

We have shown how species’ feeding strategies and the topology of the avian component of food webs differ between protected and unprotected areas across diverse biogeographical regions of Europe. We have identified potential drivers of these contrasts. We found that more than half of the European PAs investigated show a lack of contrast in food web structure compared to their surroundings. This suggests either a weak effect of protection on food webs or potential spill over effects from protected to unprotected areas. PAs specifically managed for the protection of birds favour intermediate species, larger top predators, and longer food chain lengths. Sites with comparatively more agricultural area inside PAs favour food webs with slightly longer MFCL, higher connectance, and more predators and links per species. The anthropogenic conditions outside PAs are also relevant for the outcome of protection, especially in describing the contrasts in MFLC and number of feeding links. Our results highlight the importance of taking a holistic approach to assessing biodiversity in general, and the effect of conservation strategies in particular. Even though we did find a contrasting effect of PAs on species richness, we were unable to tie these differences to a particular driver (see full results in Supplementary Table 1). By contrast, many abiotic features emerged as potential drivers of changes in biodiversity when ecological interactions, and their structure, are considered. Moreover, the comprehensive evaluation of natural communities adopted here, demonstrates the importance of moving beyond species to develop more accurate, multifaceted biodiversity assessments.

The large proportion of neutral contrast in food web metrics aligns with previous findings on the mixed effect of PAs on population trends (e.g.^16^). This was the case even for metrics like mean generality and vulnerability related to species’ specialisation contrary to the results of Cazalis et al. ^3^. Such lack of contrast in food web structure between protected and non-protected communities could arise from different mechanisms. Firstly, in Europe, where most of the landscape has been heavily anthropized, most PAs allow for some level of anthropogenic activities (e.g. IUCN category V) and are often designated for multiple functions, including recreational purposes^39^ or the historical protection of game species^40^. Thus, thinking of PAs as a single disturbance gradient is reductive. They can act on many different facets of our landscapes, from reducing land use intensification, human density to conserving cultural values. This multifunction and disparity in PA status across different European countries and different legislations could be responsible for the blurred signal in terms of protection outcome. Secondly, species dispersal across landscapes, and the resulting rescue or drainage effects, have been shown to promote biodiversity at regional scales in simulated heterogeneous landscapes, even in the presence of highly disturbed habitats^41^. Finally, on the contrary, homogeneous landscapes, e.g. remote or relatively untouched sites might display similar food web structure inside and outside protected areas regardless of whether some land is put aside for protection. In this case, the protection status of the land is mostly virtual and is independent of habitat quality or anthropogenic disturbances across the landscape.

### The importance of management practices

We found that PAs managed for birds (PAs under the Natura 2000 bird directive and the Ramsar sites for wetlands) are more effective at hosting more intermediate bird species longer MFCL and larger top species. Thus, those two conservation strategies that date back to the Bern convention in 1979 when the Natura 2000 sites and the Birds Directive were created to “support the provision of a coherent EU-wide ecological network of sites”^40^ seem to significantly enhance some aspects of food webs, attracting larger bodies predators and promoting longer food chains.

Agricultural areas seem strongly coupled with simpler food webs, but also slightly longer MFCLs. PAs on average had less agricultural area than their surroundings, and only four protected areas displayed positive difference in agricultural area (i.e. more agricultural land inside reserves). Overall, PAs hosted half as much non-irrigated arable land (large crop fields with no water reservoir nearby) as non-protected areas and slightly less pasture, in favour of forest and grassland (Supplementary Figure 2). Thus, those “higher levels of agriculture” need to be nuanced as those are still relatively small compared to levels outside protected areas. It seems that the contrast in mean food chain length might have a bell-shaped relationship with the difference in agricultural area, where the contrast in mean food chain length peaks around zero – no difference in agricultural land inside and outside protected area. This cannot be confirmed due to the scarcity of data points. Finally, given that MFCL and connectance typically increase with primary productivity^42^, PAs with higher proportion of agricultural areas may also be situated in more productive areas (which is not usually the case for PAs, which are usually relegated to areas of low economic importance). If this is right, then it suggests that moderate levels of agriculture can be compatible with some positive outcomes such as longer food chains but might also promote the simplification of food webs.

### Managing PA surroundings

Conditions outside PAs and site wide features, notably the detrimental effect of human density and urban habitat surrounding PAs were linked to the effects on MFCL and mean number of feeding links. As highlighted by previous theoretical work on the design of PAs^43^, the integrity of the landscape outside of reserves can promote species persistence inside reserves. This importance of surrounding habitats increasingly considered in the design of reserves, with the integration of buffer zones and dispersal corridors now usually implemented as “lower category” protected areas ^13^. Site-wide metrics including remoteness and land cover diversity were predictors of contrast in mean food chain length and contrast in mass of top species. The most remote PAs in our study were those in Scandinavia. In those areas, the concept of reserves may be mostly virtual, as habitats are likely rather untouched regardless of the protection status. Nonetheless, the most remote sites seemed to host reserves with shorter food chains compared to their surroundings, suggesting that habitats in those reserves are less suitable to host many intermediate and high trophic levels with many feeding links, maybe due to their lower productivity or adverse climatic conditions. The importance of the diversity of land cover types for conserving larger top predators is interesting and can be linked to their vulnerability to habitat fragmentation ^44^ and their natural occupation of larger areas ^33^.

### Data limitations

Although our original aim was to make our study as global as possible in terms of species, geographical coverage and types of PAs, scarce biodiversity survey data caused our study to be biased towards specific European bioregions (Atlantic, Alpine and Mediterranean), and taxonomic groups (bird species only). Indeed, 94.5% of the European occurrence records downloaded from GBIF were for avian species, with only 4.1% of mammals and 0.9% amphibians and 0.5% reptiles. This highlights the important limitations caused by this well-known taxonomical bias towards avian taxa in citizen science data^45^. Given that many PAs do not have as primary focus the protection of birds, but aim at preserving other species or aspects of ecosystems ^46^, it would be relevant to repeat this analysis on the full food web of tetrapods, perhaps using broader range maps (similarly to e.g. ^42^) instead of occurrence data, in order to compare results. This will represent a trade-off between data resolution / confidence and spatial scope, but could provide further insights into the conservation of food webs. Here, we opted for better local resolution / data confidence, in order to be as accurate as possible in quantifying species presence and community composition. Another limiting factor precluding a more comprehensive analysis of food webs in PAs across the globe is the availability of ecological interactions data. Even though several ecological networks have been characterised across the globe (e.g. web of life www.web-of-life.es, GloBI www.globalbioticinteractions.org, or Mangal https://mangal.io/), to our knowledge, the Tetra EU^31^European tetrapod food web is the only well-resolved continental-scale ecological network characterised to date. Filling this data gap is fundamental to develop a comprehensive assessment and understanding of the effectiveness of PAs at preserving a fundamental aspect of biodiversity: the structure of ecological interactions.

Neutral contrasts in food web metric do not inform on the food web’s absolute state (good or bad). Thus, measuring the contrast in food web metrics can be ambivalent: a positive contrast can safely be attributed to the protected communities being different from their surrounding communities, while a null difference between two healthy values in food web metrics could also be a positive outcome, suggesting that PAs and their surroundings benefit from each other in a positive feedback loop. The relatively small proportion of positive outcomes found here does not suggest that protection is useless. This nuance is important and is the reason why we refrain from talking about PA performance or effectiveness, given that our comparison lies elsewhere. We simply aim to uncover how food web structure varies spatially with respect to PAs, rather than quantifying the effect of protection (the “legal” act of assigning a protection status to an area), which would require more data and stricter filtering as well as a before after comparison (see e.g. ^16^), complementing to the spatial comparison.

Our results beg a few questions. First, it is not straightforward to identify which food web metrics are to be preserved. Depending on the conservation goal, we might wish for longer food chains, to promote trophic specialists species, or to conserve keystones species ^47^ or more connected species to maintain the integrity of food webs in case of disturbances. As our results suggest, the importance of clear conservation goals as was done for the Birds Directive and the Ramsar sites seem to be important in promoting positive outcome for communities.

Our results support the creation of reserves with favourable context and well set management goals, following the positive effects of the proportion of protected areas managed for birds and the importance of landcover type and anthropogenic pressure inside and outside reserves. Had we only considered species richness in our study, we would have concluded that no drivers had any influence on contrasts in food web metrics. Our results provide insights into the benefits of evaluating the outcome of protection from a community perspective and feed into the growing realisation of the importance of surrounding landscapes for PAs.

## Methods

### Protected areas

The data on protected areas was extracted from the World Database of Protected Areas (WDPA)^34^. The WDPA was processed by removing UNESCO world heritage sites and maritime PA, transforming all multi-polygons into simple polygons and removing polygons with an area smaller than our grain size – 10km² (as in ^48^) and PAs designated after 2015.

### Trophic interactions and species occurrence

We focused our study on species included in the tetrapod European food web Tetra-EU^31^, a species-level trophic (i.e. comprising feeding, predator-prey, interactions) metaweb comprising 1,151 tetrapod species and 60,435 trophic interactions, because to our knowledge, it is the only well-resolved continental-scale ecological network characterised to date. This limited the geographical scope of our study to the European continent. We used species occurrence data downloaded from two citizen science projects: Global Biodiversity Information Facility^29^ and eBird^30^.

About a third of the occurrence records came from the May-2022 version of eBird for all European countries except Russia. The eBird data was filtered (see Supplementary Methods for details) keeping records of species of birds included in the European food web for the breeding period (April to August). The GBIF data was downloaded using the *rgbif* package (see supp for dataset DOIs), filtering for human observations recorded after 2000, with spatial coordinates, no major geospatial issues and with coordinate uncertainty smaller than 10 km. Post-download, we further filtered the data for species-level observations again, for the breeding period (April to August) and removed presumed negated or swapped coordinates flagged by GBIF.

To homogenise the taxonomy between GBIF, ebird and the European food web we used the *taxize* package in R^49^ to match species’ scientific names. We then manually matched the species that remained unmatched using synonyms from Avibase. This resulted in 509 bird species considered for analysis.

### Study design

#### Comparison of protected and surrounding non-protected grid cells

We divided Europe into 10×10 km grid cells. Each grid cell was assigned one (or several) protection status by intersecting the WDPA shapefile with the European grid. For grid cells intersecting with several overlapping PAs, the grid cell was assigned the higher protection level (scored from of 1 to 10, from no status recorded, regional parks to national park or strict nature reserves). For overlapping PAs with identical protection levels, we filtered by age, keeping the oldest. Finally for PAs with identical age and protection level, we kept the largest. For all the remaining overlapping PAs, we randomly assigned one protected area to the grid cell.

We calculated sampling effort as the number of survey events per grid cell throughout the study period. One survey event was defined as a unique survey visit to a location: the combination of date, time, location, and survey program from which the record originated (e.g. INaturalist, observation.org, eBird etc). We removed all grid cells with a number of survey events under 50 survey events. Additionally, we rarefied the number of survey events per grid cells N to that of the grid cell with smallest number of survey events Nmin of each site to ensure communities were comparable within sites. This was done by randomly sampling without replacement Nmin survey events from the pool of N survey events from each grid cell.

Hereafter, we differentiate the “whole dataset” from the “subset dataset”. The whole dataset is the dataset with all the grid cells before filtering for survey effort. It was used to characterise the sites in terms of environmental characteristics (see below). The subset dataset is the dataset after filtering for survey effort. This dichotomy is illustrated in Figure 1.

#### Grouping PAs into sites

To account for the fact that, in Europe, PAs are often clustered and small, we proceeded to group “connected” PAs (i.e. within 1km of each other). In addition to ensuring higher number of data points, we deemed this method more realistic in terms of the scale at which birds tend to move, as well as a way to account for the complementarity and non-independence of different types of closely situated protected areas ^50^. To compare protected grid cells to their non-protected counterparts in similar environmental conditions, we restricted our comparison to the cells within 100 kilometres of a PA and in the same bio-geographical region (see below). A set of protected and its non-protected grid cells is hereafter called a site.

Our use of 100 km buffer zone is a large area to consider as comparison. This large number was chosen mostly to increase our sample size of sites considered. In practice however, protected areas being so clustered in Europe, 75% of our sites were compared with unprotected communities within 55 kilometres of their boundaries.

For each site, we kept only occurrence records that were recorded after the designation of the youngest PA within each site. For example, if the site is constituted of 10 PAs among which the youngest PA was designated in 2010, we keep occurrence records from 2010 onward (12 years of records) for the whole site – to ensure that all our local communities were protected at the time when their bird population was surveyed. In addition, we only kept sites with at least 15 protected grid cells and 15 non-protected grid cells yielding a total of 46 sites.

From these 46 sites, one very large site situated in Poland had very few sampled communities relatively to its total area (a ratio of 3% of the communities were correctly sampled, all other sites being between 12 and 97%). Thus, we deemed this site to be poorly represented by our occurrence data and removed it from all subsequent analysis. Thus, our analysis was conducted on 4591 communities (i.e. grid cells) spread across 45 sites distributed mostly in Western Europe (Spain, France, Great Britain) with some sites also in Germany, Poland, Austria, Norway, Sweden and Bulgaria (Figure 1). The average site was made of 7 (sd = 6, min = 1, max = 123) protected areas. In the whole dataset (all grid cells, irrespective of survey effort – transparent grid cells in Figure 1), sites had on average 66 protected grid cells (min = 16, max = 397) and 132 non-protected grid cells (min = 20, max = 543), whilst in the subset dataset with only well surveyed cells (opaque cells in Figure 1), sites had on average 31 protected grid cells (min = 15, max = 99) and 70 non-protected grid cells (min = 19, max = 348).

### Food web metrics

Thirteen topological network properties were calculated on the food webs constructed locally at each grid cell (dataset with food web metrics can be found in Supplementary Data 2). We chose metrics that measured community complexity, topology and described the identity of their species, and that have been hypothesised as important drivers of food web stability. These included measure of the topology of the food web, such as their compartmentalisation, mean food chain length and fraction of omnivory as well as its complexity, with measure of the number of links, fraction of realised links as well as number of links per species. Finally, we included measures of species composition with the number of basal, intermediate, and top species, as well as their body size. More details on the definition of those metrics can be found in Supplementary Methods.

Given that our study is concerned only with birds, the notion of basal species in our studies refers to birds that don’t feed on other birds (insectivorous, granivorous etc) as opposed to the “classic” idea of a basal species that are usually either herbivores or primary producers.

### Statistical Analyses

To conduct a robust comparison between protected and non-protected grid cells we computed the contrast in network metrics between protected and surrounding non-protected cells (*Within-site contrast in food web structure between protected and non-protected communities*) by controlling for biases in PA location, then we assessed what drivers best explained those differences (*Drivers of protection effectiveness across sites*) (Fig. 5).

**Fig. 5.**
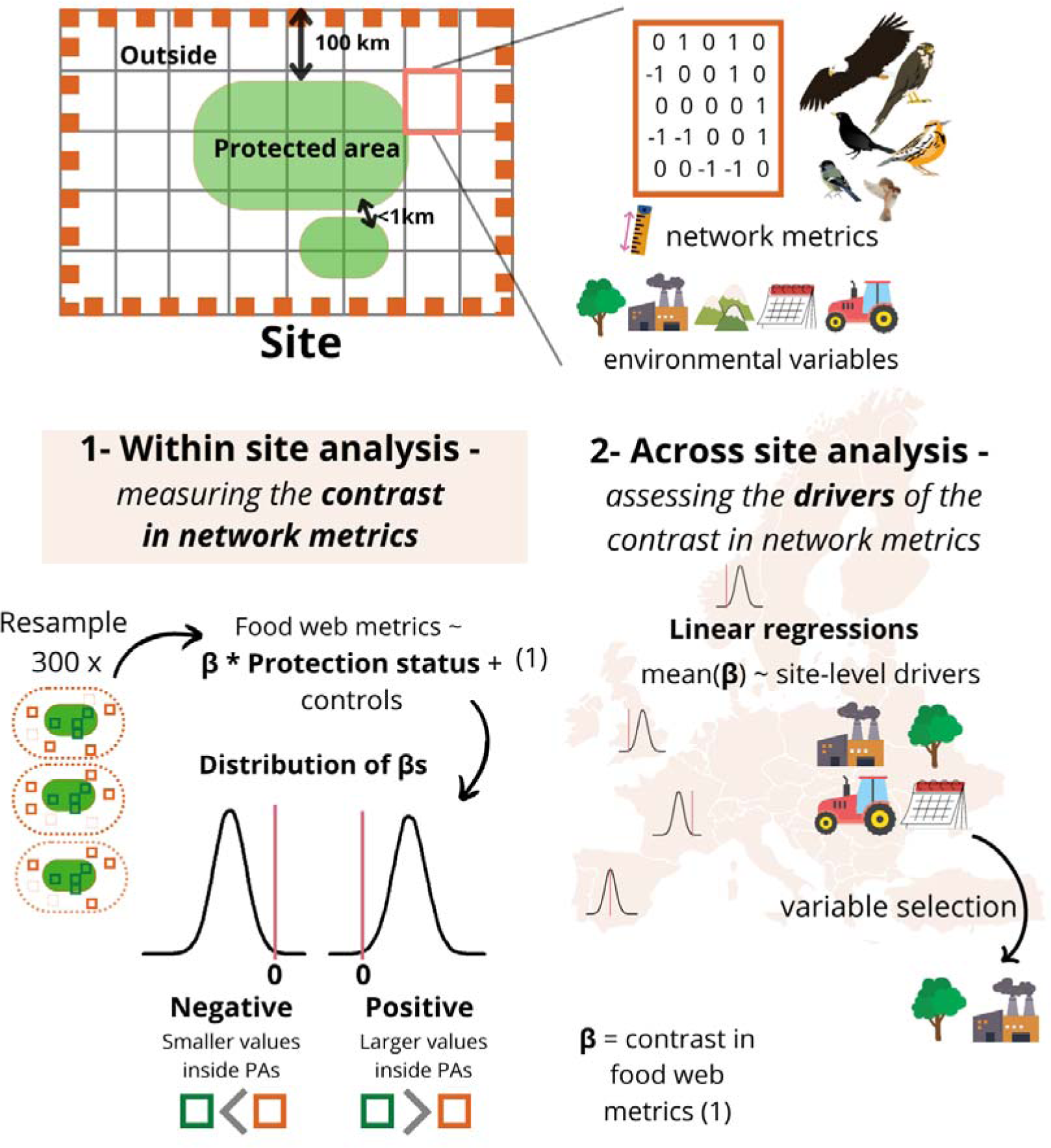
Summary of the three main stages of analysis. The three panels correspond to one subsection to our framework. (1) First, we constructed 45 sites made up of connected protected areas (distant from less than 1 kilometre) and their surrounding non-protected communities (within 100 kilometres) across Western and Scandinavian Europe. Each site was divided into 10×10 kilometre grid cells, which each held a community of interacting birds. For each community, we computed its food web structure and environmental conditions. (2) The second step of our analysis was to quantify the within-site contrast in food web metrics between protected and surrounding non-protected communities. This was done by running 45 (sites) x 13 (metrics) Generalised Additive Mixed Models (GAMMs) on 300 bootstrapped samples of local communities to ensure a balanced number of protected and non-protected communities. The contrast in network metric corresponded to the coefficient for binary protection status in our models. Thus, for each site we obtain a distribution of 300 contrasts in food web metrics (s). We consider that the difference in food web structure was significant if the 95% quantiles of these distributions did not overlap zero. (3) The last step of our analysis was to quantify the role of the environmental context of the sites as an across-site predictor of protection outcome. To quantify this, we ran simple linear models of the mean contrast in food web metrics (mean *β*) against a suite of environmental predictors, including land cover type, human pressure and protected area characteristics.

#### Within-site contrast in food web structure between protected and non-protected communities

For each site, we quantified the contrast in food web structure between protected and non-protected food webs. This was conducted using Generalised Additive Mixed Models (GAMMs) (*mgcv* package^51^).

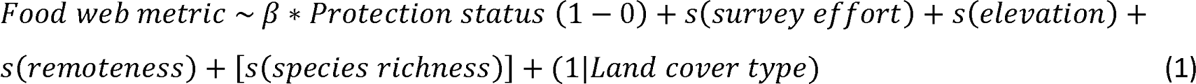

Here, the *β* coefficient (1) for the binary (1/0) “protection status” term in our model corresponds to the contrast in food web metrics: Identically to traditional linear regression, this coefficient corresponds to the difference in means between the “reference” group (here “1 = protected”), and that of the other group(s) (here 0 = “Non-protected”). Computing the difference in this way allowed us to statistically control for within-site differences in environmental conditions, survey effort and species richness. This was done using non-linear smoothers represented as the *s* terms in Eq. 1. Indeed, protected area location are known to be biased towards sites that are “unlikely to face land conversion even in the absence of protection”, i.e. at higher elevations, steeper slopes and greater distances to roads and cities^38^. Our control terms included a random effect for land cover type (CORINE Landcover^52^), and non-linear smoothing terms for elevation ^53^, remoteness ^54^ (see Supplementary Methods for further details), survey effort and species richness (for all food web metrics except species richness). In addition, to ensure that we are comparing sufficiently similar communities in terms of climatic conditions and species composition, we only compared grid cells within the same bio-geographical region (<www.eea.europa.eu>).

Different error distributions were used depending on the nature of the food web metric (dependant variable): count metrics were modelled using a negative binomial error distribution, proportion data using quasibinomial error distribution with a log link and continuous metrics using a gaussian distribution.

For each model (45 sites x 13 food web metrics), to ensure a balanced number of protected and non-protected grid cells, we resampled the most abundant category of grid cells (5 were inside and 40 outside) without replacement 300 times to the size of the least abundant category and thus ran 300 iterations of each GAMM. On average the GAMMs were run on 60 (sd = 33, min = 30, max = 198) grid cells (half being protected and the other half not).

This yielded a distribution of 300 *β* coefficients for the contrast in network metrics (see example in Figure 2) from which we extracted the mean and 95% quantiles to assess its direction and “significance”. Here, we consider the contrast in network metric between inside and outside to be “significant” if the 95% quantiles from the distribution of the contrasts in network metrics do not overlap zero.

We compared GAMMs with their equivalent Generalised Linear Mixed Models (GLMM) - which only allow for linear effects -, and found an average AIC difference of 58 points, in favour of GAMMs.

#### Drivers of protection effectiveness across sites

To assess the role of environmental conditions in modulating the outcomes for the contrasts in metrics, each site was characterised by 15 environmental drivers (more details can be found in Supplementary Methods):

1. *protected area characteristics* including the average date of designation of its constituting PAs, average protection level, the aggregation of its protected areas, the proportion of forested areas inside the reserves and the proportion of protected areas managed for birds or protected areas under the Natura 2000 bird directive and the Ramsar sites for wetlands ^16^.
2. *Sites level characteristics* including average elevation, slope and land cover diversity based on Corine Landcover’s coarsest classification^52^ (see Supplementary Methods for more information).
3. *characteristics of the surrounding landscape* including the mean human density ^55^, and the mean proportion of urban landscape, proportion of agricultural area and proportion of forested area (see Supplementary Methods for details on how those variables were computed).
4. the *contrast with surrounding landscape* was computed as the difference between the mean values inside and the outside for the proportion of agricultural areas, forested areas, human density and urban habitat.

This was done by relating the mean contrast in food web metrics 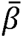 of each of the 45 sites to their environmental context using simple linear regressions, controlling for the proportion of protected grid cells that varied quite strongly across sites (Eq. 2).

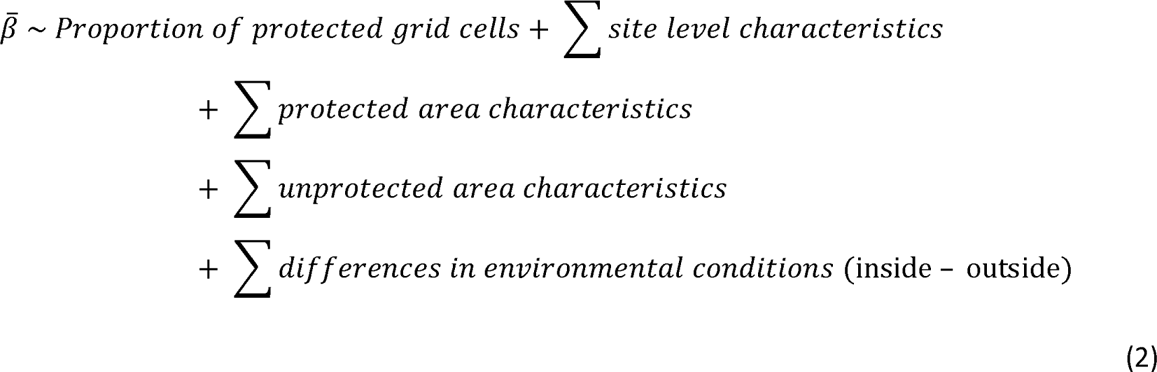

Given our small number of sites (N = 45), we used variable selection to reduce the number of predictors in our models. We compared used variable selection based on BIC. We extracted model coefficients, standard errors, p values and adjusted R² to assess the performance of the linear models and the direction and magnitude of the effect of each variable on the contrast in network metrics. Confidence intervals (at 95%) for visualisation were extracted from the multiple regression visualisation tool *visreg*^56^. We used a redundancy analysis (RDA) to illustrate the results and correlation between drivers and network metric.

All analyses were run on R version 4.3.0 ^57^.

## Data availability

Data produced for intermediate steps in the analyses (Supplementary Data) will be made available will be made available once this manuscript gets accepted in a peer-reviewed journal.

All data were extracted / downloaded from open sources. Biodiversity records are accessible from the GBIF at https://www.gbif.org/. dois for the GBIF downloads are available in Supplementary Methods – Table 1.

eBird version of May 2022 for all European countries is available at http://www.ebird.org.

Land cover data^52^ are available from <https://land.copernicus.eu/pan-european/corine-land-cover/clc2018/fetch-land-file?hash=83684d24c50f069b613e0dc8e12529b893dc172f>

The TETRA-EU 1.0 dataset of species interactions^31^ and bio-geographical region shapefile are available at <https://doi.org/10.5061/dryad.jm63xsj7b>

The World Database on Protected Areas (WDPA)^34^ March 2022 version available at <www.protectedplanet.net>

Elevation from the Shuttle Radar Topography Mission (SRTM)^58^ available at https://www.worldclim.org/data/worldclim21.html under “Elevation”

Remoteness data^54^ available at < https://malariaatlas.org/> under “Travel time to cities”

Human density data for 2015 came from the Gridded Population of the World dataset, version 4.11 (GPWv4.11, UN WPP-Adjusted Population Density, v4.11, 2015)^55^ available at < https://ghsl.jrc.ec.europa.eu/download.php?ds=pop >

## Code availability

The computer code for all the analyses (including data processing) presented in this study will be made available once this manuscript gets accepted in a peer-reviewed journal.

## Acknowledgements

LT was funded by Swansea University ECR BIOL postgraduate research scholarship. NG received funding from the European Union’s Horizon 2020 research and innovation programme under the Marie Skłodowska-Curie grant agreement BIOFOODWEB (No 101025471).

## Author contributions

M.L. and L.T. conceived the study. M.L. and L.T. designed data-related protocols and analyses with input from N.G. and K.W. L.T. collated and processed data. L.T. conducted analyses. L.T. and M.L. led manuscript preparation, and N.G. and K.W. contributed to substantive revisions and edits.

## Additional Information

**Supplementary information** associated to this manuscript is presented in an attached document.

