## Supplementary Figures for "Protected areas enhance avian food webs"

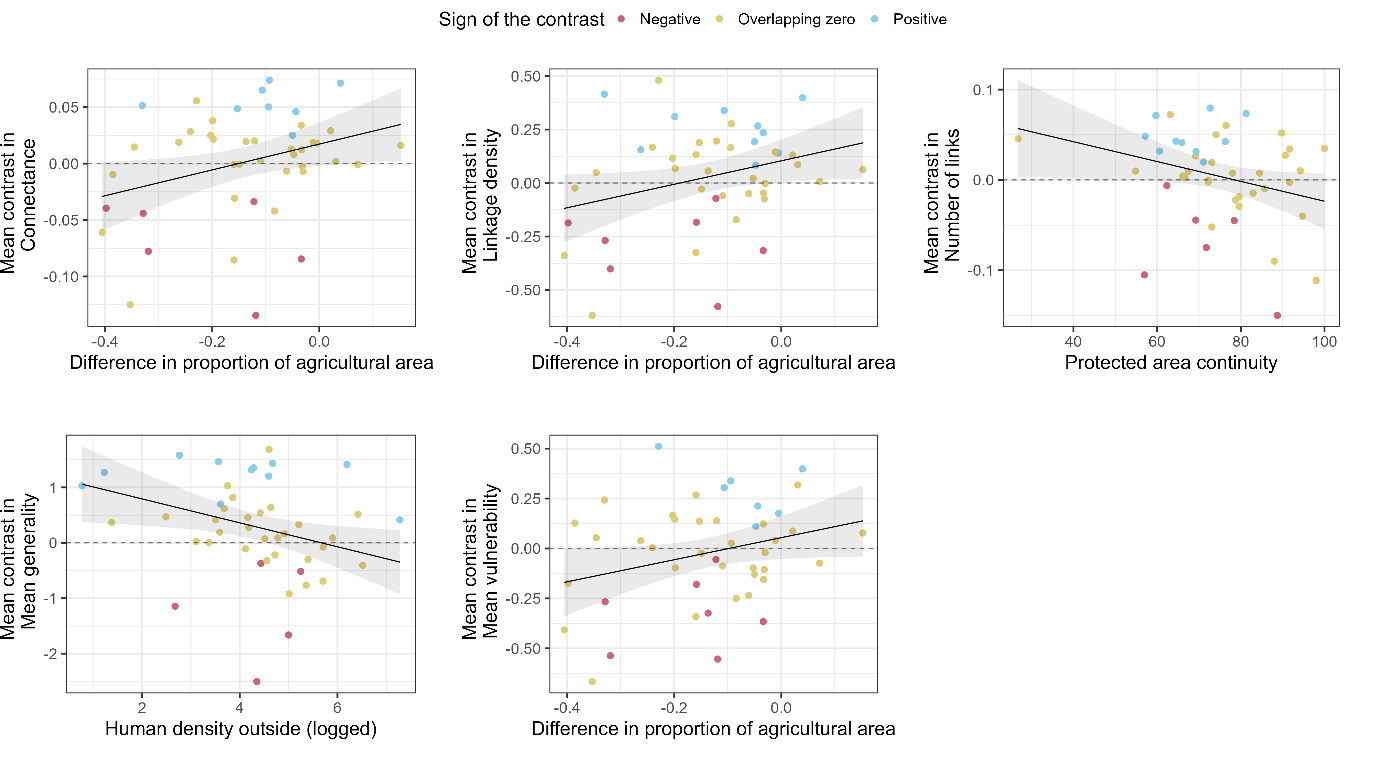

**Supplementary Figure 1.a.** **Full illustration of the partial effects of raw (non-scaled) drivers on the mean contrast in food web.** 95% confidence interval for multiple linear regressions were extracted from the “visreg” package. Results from variable selection based on BIC criterion in simple linear regressions relating the mean contrast in food web metrics β ̅ against scaled environmental drivers (Supplementary Data 2 ) across 45 sites (see Supplementary Figure 1 for non-scaled coefficients). The data used for the regression is the average differences across 300 bootstrapped iterations of randomly chosen pairs of cells inside vs outside of each PA (Fig. 1 and Methods). Each linear model also included one extra variable to statistically account for the variable proportion of protected areas across sites, as well as an intercept term (see Supplementary material 3 for the full scaled results and Supplementary Figure 4 for the non-scaled results). Drivers with a significantly positive coefficient are in blue (these drivers tend to occur in sites with positive contrast in food web metrics, i.e. larger food web metrics inside than outside protected areas – see Fig. 2 in main text) and negative coefficients are in red (these drivers occur tend to occur in sites with negative contrasts in food web metrics i.e. larger food web metrics outside than inside protected areas). The variations in contrast for other food web metrics were poorly explained by our set of drivers – none were selected by BIC step selection - which is why they are not included in this plot.

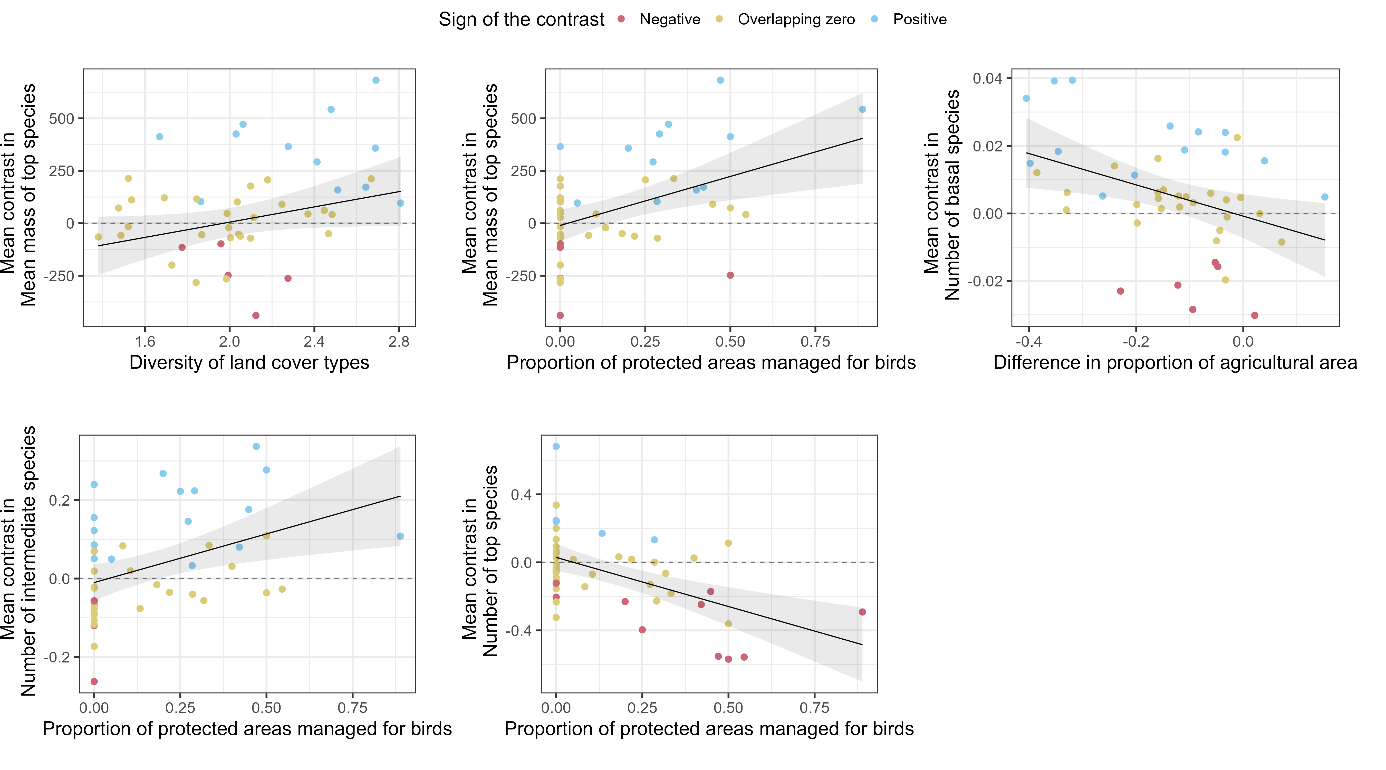

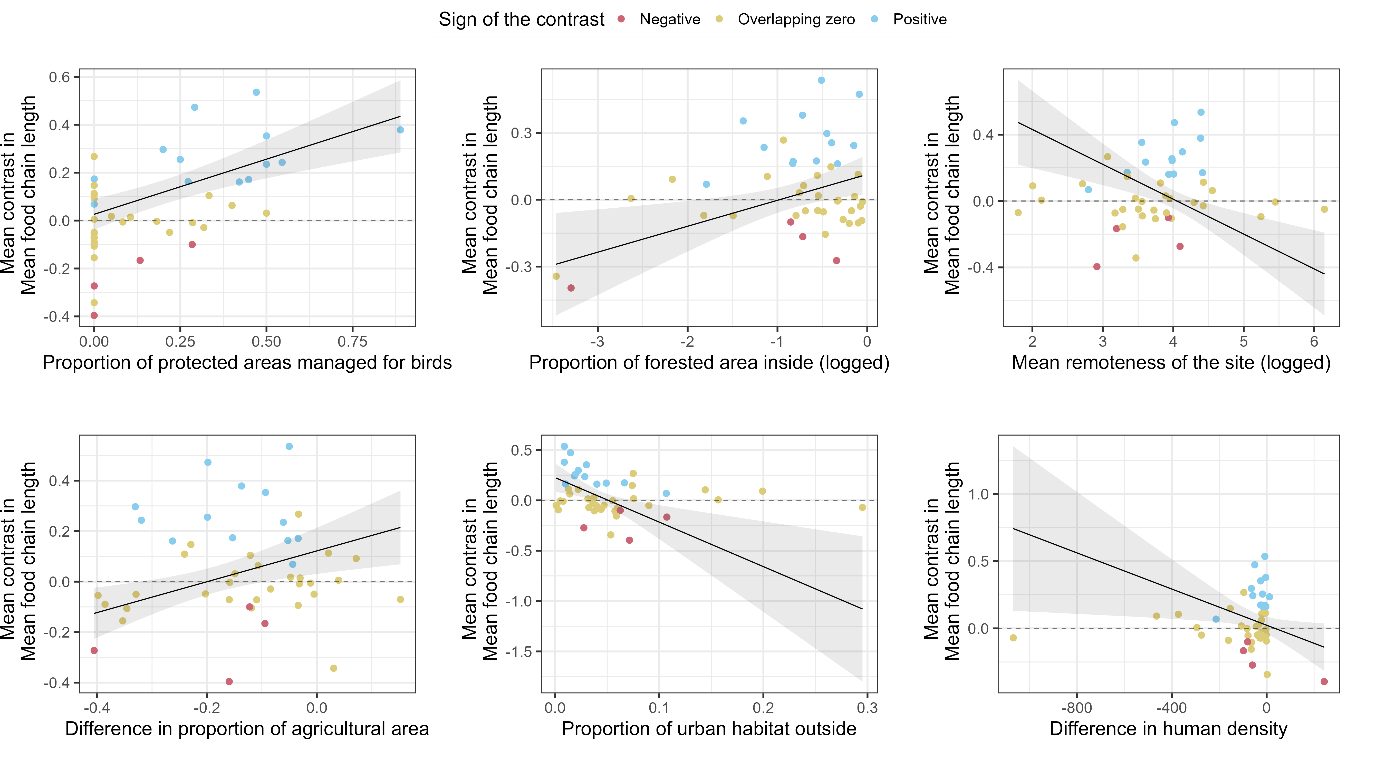

**Supplementary Figure 1.b. Full illustration of the partial effects of raw (non-scaled) drivers on the mean contrast in food web.** 95% confidence interval for multiple linear regressions were extracted from the “visreg” package. Results from variable selection based on BIC criterion in simple linear regressions relating the mean contrast in food web metrics β ̅ against scaled environmental drivers (Supplementary Data 2 ) across 45 sites (see Supplementary Figure 1 for non-scaled coefficients). The data used for the regression is the average differences across 300 bootstrapped iterations of randomly chosen pairs of cells inside vs outside of each PA (Fig. 1 and Methods). Each linear model also included one extra variable to statistically account for the variable proportion of protected areas across sites, as well as an intercept term (see Supplementary material 3 for the full scaled results and Supplementary Figure 4 for the non-scaled results). Drivers with a significantly positive coefficient are in blue (these drivers tend to occur in sites with positive contrast in food web metrics, i.e. larger food web metrics inside than outside protected areas – see Fig. 2 in main text) and negative coefficients are in red (these drivers occur tend to occur in sites with negative contrasts in food web metrics i.e. larger food web metrics outside than inside protected areas). The variations in contrast for other food web metrics were poorly explained by our set of drivers – none were selected by BIC step selection - which is why they are not included in this plot.

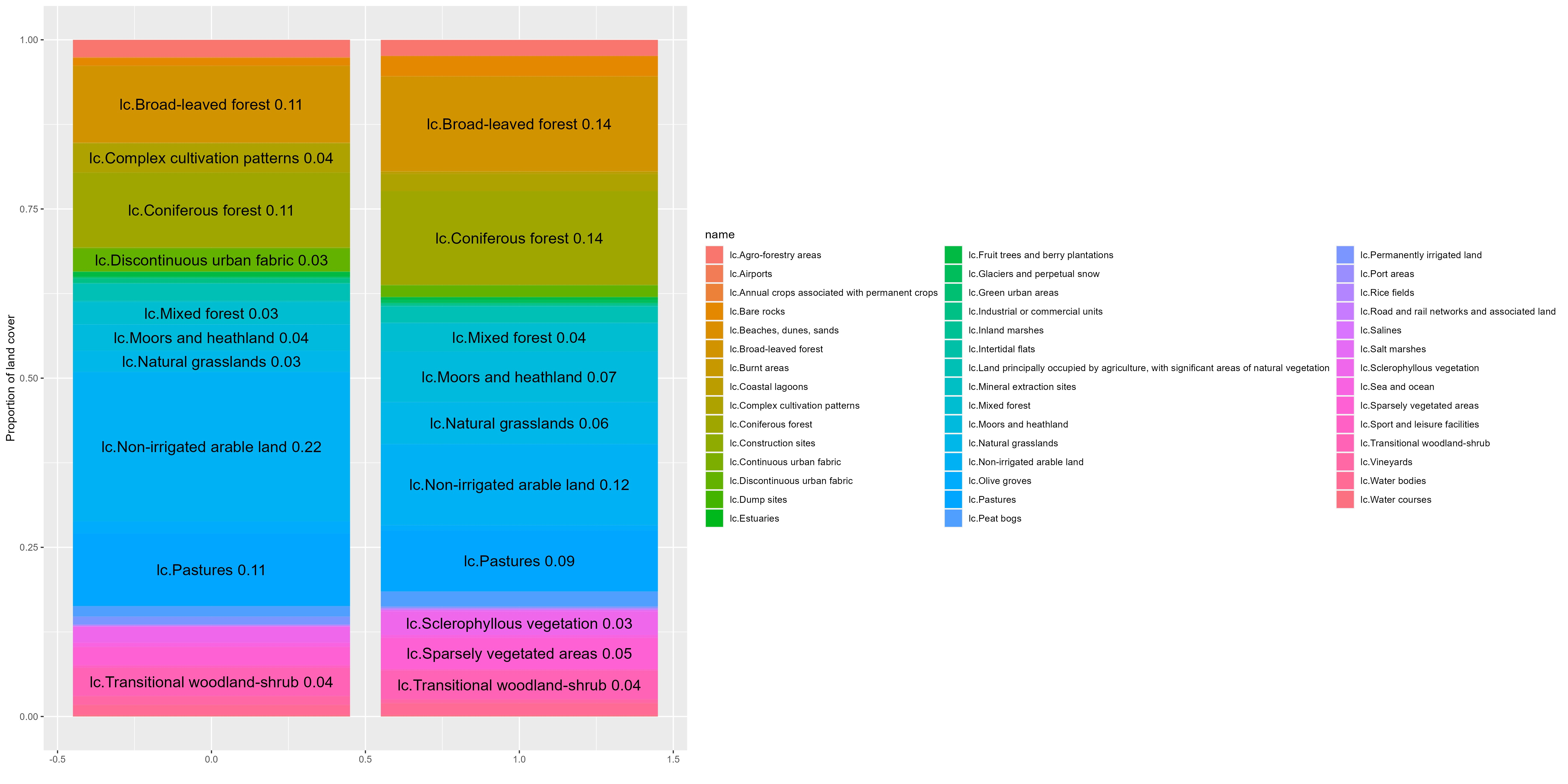

**Supplementary Figure 2. Differences in land cover between non-protected communities (left) and protected communities (right).** Only proportions larger than 0.03 are printed. Land cover categories are from the finest classification of corine land cover dataset (Copernicus Land Monitoring Service & European Environment Agency, 2018).
