## Supplementary Methods for "Protected areas enhance avian food webs"

### Protected areas

#### Protected area data filtering

Protected area shapefiles (WDPA) were downloaded from www.protectedplanet.net/. The dataset is split into 3 shapefiles that all span the globe. The WDPA was processed in the R script 2.FilteringWDPA.R.
We first removed predominantly or entirely marine protected areas, UNESCO-MAB Biosphere Reserves (following^1^), protected areas whose implementation was not complete, thus keeping only status "Designated", "Established" or "Inscribed". To fit with our chosen occurrence data resolution of 10km², we removed any protected area with a GIS area smaller than 10km², and subset the dataset to the extent covered by the European interaction database^2^ (Supplementary Methods Fig. 1).

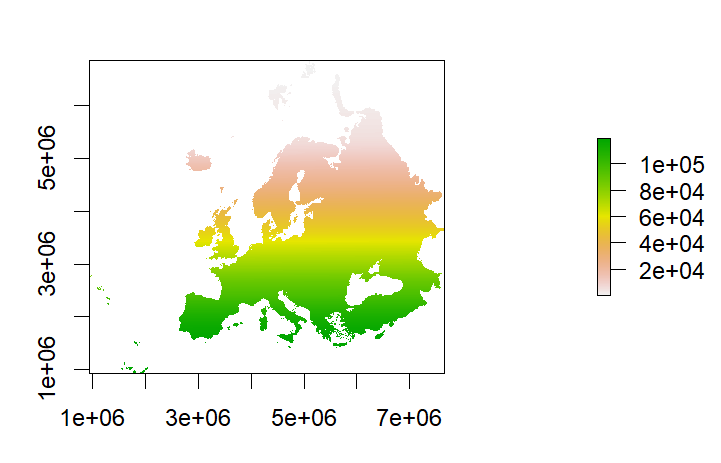

Supplementary Methods Fig. 1 : Spatial extent of the European interaction database^2^ TETRA EU 1.0. Grid cells are coloured by ID.

Because in the WDPA protected areas can be constituted of several individual polygons under the same WDPAID, we split those multi-polygons into simple polygons and removed any remaining polygons smaller than the 10km² resolution.

Finally, we removed protected areas designated after 2015 to ensure that all protected areas would have at least 7 years of occurrence data to print an accurate picture of their current community composition.

The filtered protected area dataset was then split into grid cells. We removed overlapping designation (e.g. the same grid cell was both a Ramsar sites (wetlands of international importance) and UNESCO World Heritage Sites) by keeping (1) the highest level of protection or if all equal (2) the oldest designation or if all equal (3) the largest polygon. If all of those three criteria were still equal, we randomly picked one designation to assign to the remaining unlabelled grid cells.

#### Grouping of Protected Areas into “sites”

We inferred the distance between neighbouring PAs by calculating the minimum distance between grid cells inside a PA to grid cells inside other PAs within 200kms. This gave an approximate value of the distance between neighbouring PAs, which was then used to group PAs less than 1km apart together into sites. Each group of PAs was then assigned a new ID and split by bioregion.

### GBIF

#### GBIF download and filtering protocol

Occurrence records from GBIF were downloaded using the rgbif package that works through the GBIF API. The code to reproduce the download is in R script 1.DownloadGBIF.R. We automated the download of the desired records, to select only species present in the interaction network database^2^, filtering for human observations recorded after 2000, with spatial coordinates, no major geospatial issues and with coordinate uncertainty smaller than 10 km. We split Europe into 9 rectangles (Supplementary Methods Fig. 2), which means that the downloaded records were split into 9 files. We initially downloaded data for all taxa in the hope to have enough records to run the analysis on all tetrapods, but ended up filtering the data for birds only for lack of data for amphibians, mammals and reptiles.

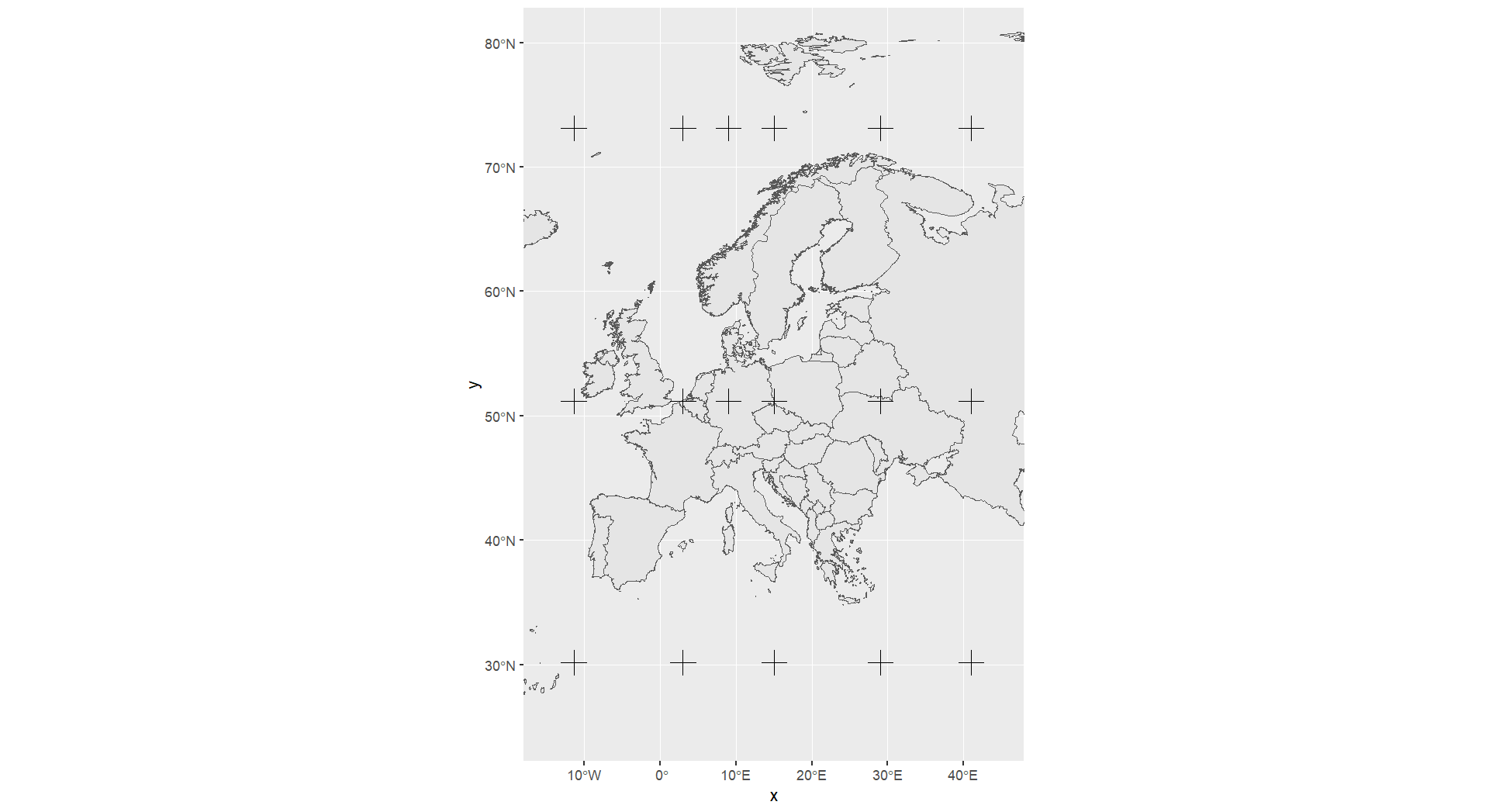

**Supplementary Methods Figure 2.** coordinates of the polygons used to split Europe into 9 rectangles for GBIF download. Each rectangle corresponds to one row in Supplementary Methods Table 1.

**Supplementary Methods Table 1. Details on the GBIF occurrence data downloads.** GBIF data was downloaded in sections to reduce the size of individual files. We divided Europe into 9 rectangle polygons (Supplementary Methods Fig. 2) and inputted the polygon coordinates in the “rgbif” package (see r code available from main manuscript). Each of the 9 dataset has a doi (citation column) for reproducibility.

| Position of polygon | Nb of occurrences | Nb of datasets | Polygon coordinates to input into GBIF | Citation |
| --- | --- | --- | --- | --- |
| North 1 | 9,792,753 | 371 | POLYGON((-11.42578 51.1946,3.00000 51.1946,3.00000 73.13515,-11.42578 73.13515,-11.42578 51.1946)) | GBIF.org (14 April 2022) GBIF Occurrence Download https://doi.org/10.15468/dl.8qxh5t |
| North 2 | 41,282,352 | 62 | POLYGON((15.0000 51.1946,29.03027 51.1946,29.03027 73.13515,15.0000 73.13515,15.0000 51.1946)) | GBIF.org (14 April 2022) GBIF Occurrence Download https://doi.org/10.15468/dl.3yfv6f |
| North 3 | 44,833,015 | 60 | POLYGON((3.00000 51.1946,9.0000 51.1946,9.0000 73.13515,3.00000 73.13515,3.00000 51.1946)) | GBIF.org (14 April 2022) GBIF Occurrence Download https://doi.org/10.15468/dl.58vda8 |
| North 4 | 55,818,583 | 45 | POLYGON((9.00000 51.1946,15.0000 51.1946,15.0000 73.13515,9.00000 73.13515,9.00000 51.1946)) | GBIF.org (14 April 2022) GBIF Occurrence Download https://doi.org/10.15468/dl.dre5hg |
| North 5 | 1,278,318 | 57 | POLYGON((29.03027 51.1946,41 51.1946,41 73.13515,29.03027 73.13515,29.03027 51.1946)) | GBIF.org (16 May 2022) GBIF Occurrence Download https://doi.org/10.15468/dl.wa8st3 |
| South 1 | 9,321,985 | 2,025 | POLYGON((-11.42578 30.17552,3.00000 30.17552,3.00000 51.1946,-11.42578 51.1946,-11.42578 30.17552)) | GBIF.org (14 April 2022) GBIF Occurrence Download https://doi.org/10.15468/dl.3n67a3 |
| South 2 | 23,486,246 | 754 | POLYGON((3.00000 30.17552,15.0000 30.17552,15.0000 51.1946,3.00000 51.1946,3.00000 30.17552)) | GBIF.org (14 April 2022) GBIF Occurrence Download https://doi.org/10.15468/dl.jchhuq |
| South 3 | 629,279 | 33 | POLYGON((15.0000 30.17552,29.03027 30.17552,29.03027 51.1946,15.0000 51.1946,15.0000 30.17552)) | GBIF.org (14 April 2022) GBIF Occurrence Download https://doi.org/10.15468/dl.yq8dhv |
| South 4 | 110,806 | 17 | POLYGON((29.03027 42,41 42,41 51.1946,29.03027 51.1946,29.03027 42)) | GBIF.org (16 May 2022) GBIF Occurrence Download https://doi.org/10.15468/dl.j5m26c |

#### Post download processing

Post-download (R script 4.DatasetCreation.R), we further filtered the data for species-level observations and non-empty occurrenceID. We removed presumed negated or swapped coordinates flagged by GBIF and records with a coordinate uncertainty larger than 10 kilometres. We then subset GBIF occurrence records to those falling within the European interaction database spatial extent^2^ (Supplementary Methods Fig. 1).

### eBird

#### eBird download and filtering protocol

About a third of the occurrence records came from the May-2022 version of eBird for all European countries except Russia. Data was downloaded from <https://eBird.org>. Files for Spain and Great Britain were particularly big and were split into smaller files.

Filtering of eBird data was heavily inspired by^3^. We kept only checklists where all species were recorded and records that were approved, and selected protocols that were either "Traveling", "Stationary" or "Historical". Only checklist that lastest at least 30 minutes and less than 600 minutes were kept. Non-stationary checklists with null distance recorded were removed and very long travelling distances (>5km) were also removed. Domestic species were removed.

The eBird data was finally filtered keeping records of species of birds included in the European food web for the breeding period (April to August) and within the spatial extent of the European food web (Supplementary Methods Fig. 1).

### Merging eBird and GBIF

#### Taxonomic homogenisation between eBird, GBIF and the European trophic interaction database

To homogenise the taxonomy between GBIF, ebird and the European trophic interaction database we implemented the taxonomic matching in^4^ using the *taxize* package to match species’ scientific names between the different datasets, using the European trophic interaction database as a reference.

We then manually matched the remained unmatched species using synonyms from Avibase at <https://avibase.bsc-eoc.org/>. This resulted in 509 bird species considered for analysis.

#### Filtering GBIF and eBird on survey effort

For both GBIF and eBird we generated a list of survey events in order to calculate survey effort. One survey event corresponded to a unique visit to a grid cell. It had a unique dataset ID (the program from which the observations originated – this could be a regional survey program or a citizen science app such as iNaturalist or eBird for example), date and time and grid cell location. A survey event was associated with a checklist of species (suite of bird occurrence records). We kept only grid cells with at least 50 survey events.

#### Integrating PA information to GBIF and eBird occurrence records

We then matched each grid cell to a group of PAs based on their grid-cell location. We only kept grid cells inside and within 100 kilometres of a group of PA. A group of PAs plus their surrounding grid cells constituted what we hereafter called a ‘site’ (Supplementary Methods Fig. 3). Thus, each survey event was assigned a site. For each site, we kept only occurrence records that were recorded after the designation of the youngest PA within each site. For example, if the site was constituted of a group of 10 PAs among which the youngest PA was designated in 2010, we keep occurrence records from 2010 onward (12 years of records) for the whole site – to ensure that all our local communities were protected at the time when their bird population was surveyed.

In addition, we only kept sites with at least 15 protected grid cells and 15 non-protected grid cells yielding a total of 46 sites.

From these 46 sites, one very large site situated in Poland had very few sampled communities relatively to its total area (a ratio of 3% of the communities were correctly sampled, all other sites being between 12 and 97%). Thus, we deemed this site to be poorly represented by our occurrence data and removed it from all subsequent analysis.

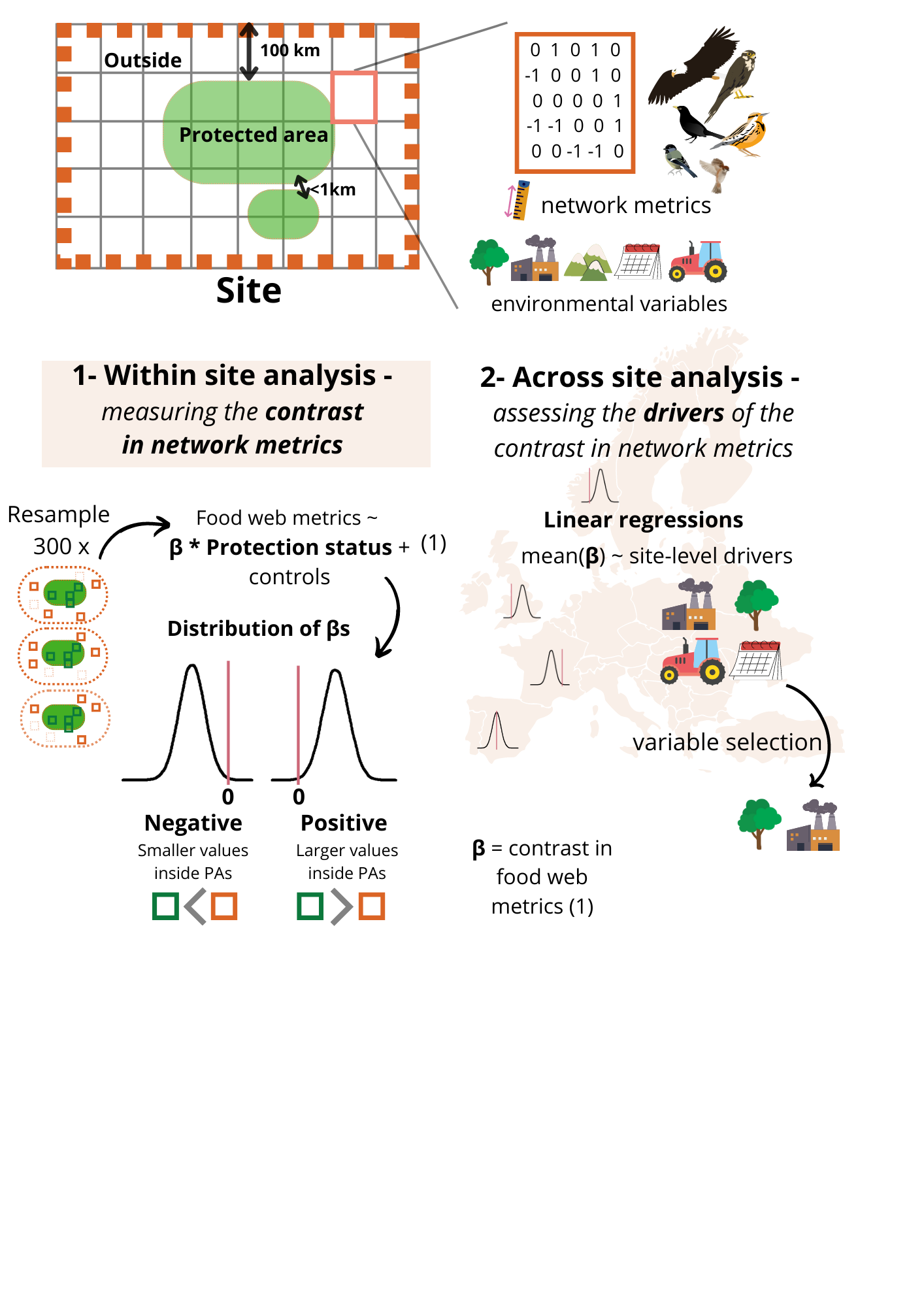

Supplementary Methods Fig. 3 : Schematic of a site, which is constituted of a group of PAs within 1km of each other and grid cells within 100 km of that group of PAs.

#### Calculation of food web metrics –

We then collated checklist of local communities, i.e. species co-occurring in each grid cell kept for analysis. We extracted trophic interactions for each one of those local communities from the European trophic interaction database^2^ and computed 13 food web metrics described in Supplementary Methods Table. 2.

**Supplementary Methods Table 2. Description of 13 food web metrics and their formulas used across the analysis.**

| Category | Food web metric name | Metric symbol | Definition | Formula | R package (when relevant) | Ecological interest |
| --- | --- | --- | --- | --- | --- | --- |
| Community-level descriptors | **Species richness** | S | Number of interacting nodes (species) in the community |  |  | Measure of the complexity of the food web |
|  | **Number of links** | L | Number of feeding links in the community |  |  | Measure of the complexity of the food web |
|  | **Connectance** | C | Proportion of realised feeding links relative to all possible ones | L/S^2 |  | Measure of the complexity of the food web |
|  | **Linkage density** | L.S | Number of links per species | L/S |  | Measure of the degree of specialisation |
| Average species’ traits in the community | **Mean vulnerability/Outdegree** | Vul  SDVul | Average number of outgoing links per nodes |  |  | Measure of the predation pressure |
|  | **Mean generality/Indegree** | Gen  SDGen | Average number of incoming links per nodes |  |  | Measure of diet breadth, degree of generalisation |
|  | **Omnivory** |  | Fraction of species with indegree larger than two and non-integer trophic level |  | Cheddar | Feeding on resources across trophic levels – measure of stability of the food web |
|  | **Fraction of basal species** | F basal | Fraction of species with no incoming links (resources) – note that here, as we consider only the bird food web so basal species are birds that don’t feed on other birds (granivorous, insectivorous, frugivorous etc) |  |  |  |
|  | **Fraction of top species** | F top | Fraction of species with no incoming links (predators) |  |  |  |
|  | **Fraction of intermediate species** | F int | Fraction of species with both incoming and outgoing links |  |  |  |
|  | **Mean chain averaged trophic level** | TL | Mean average number of nodes between the basal species and each node |  | *Cheddar/netcarto* |  |
|  | **Mean food chain length** | mfcl | Mean trophic chain length (mean number of nodes between basal species and top predators) | sum(trophic chain length – number of nodes between each basal species and each top species)/number of chains | *Cheddar* | Correlates with habitat quality – better resources enable longer food chains |
| Architecture of the network | **Modularity** | M | Maximisation of modularity function - difference between within-module (compartments) links and between-module links where a module is a subset of the network | $\sum_{s=1}^{N_{m}} \left[ \frac{L_{s}}{L}-\left( \frac{d_{s}}{2L} \right)^{2} \right]$ with $N_{m}$the number modules, $L_{s}$the number of links in module *S*, and $d_{s}$ the sum of degree of module *S* | *iGraph* | Measure of food web complexity – related to food web stability |

### Environmental data

#### Within site analysis -

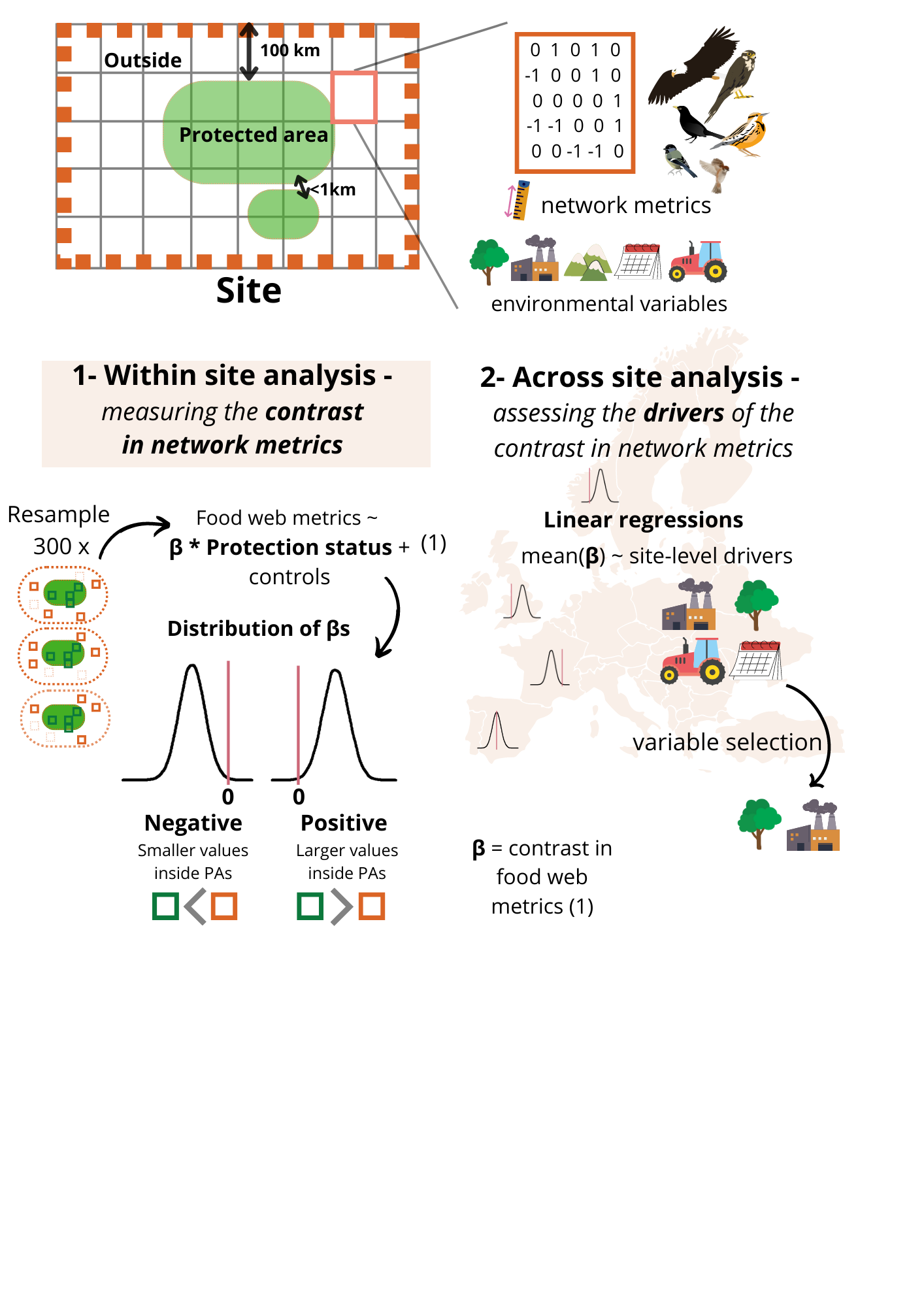

Supplementary Methods Fig. 4: The second step of our analysis was to quantify the within-site contrast in food web metrics between protected and surrounding non-protected communities. This was done by running 45 (sites) x 13 (metrics) Generalised Additive Mixed Models (GAMMs) on 300 bootstrapped samples of local communities to ensure a balanced number of protected and non-protected communities. The contrast in network metric corresponded to the β coefficient for binary protection status in our models. Thus, for each site we obtain a distribution of 300 contrasts in food web metrics (βs). We consider that the difference in food web structure was significant if the 95% quantiles of these distributions did not overlap zero.

Bioregion, land cover type, elevation and remoteness were all integrated in the food web metric contrast calculation (see Supplementary Methods Fig. 4 and main text for more information on analysis) to control for biases in PA location^6^.

##### a.1- Bioregions

The European Environmental Agency (EEA) has defined a zonation of Europe into biogeographical regions (hereafter bioregions), based on similarities in environmental conditions and habitats. As in^7^ we used 10 European bioregions from this classification: Alpine, Anatolian, Arctic, Atlantic, Black Sea, Boreal, Continental, Mediterranean, Pannonian and Steppic. A full description of each bioregion is available online (<www.eea.europa.eu>). A bioregion was assigned to each site.

Only cells within the same bioregion were compared.

##### a.2- Land cover type

Land cover data was extracted from the CLC 2018 version v.2020_20u1 raster which classified land cover into 44 classes (on level-3) and five main land cover groups: Artificial surfaces, Agriculture, Forests and seminatural areas, Wetlands and Water; for 25 hectare minimum mapping unit and 100 meter minimum mapping width^8^. For each grid cell we extracted the proportion of each broad land cover type (agricultural areas, artificial surfaces, forest and semi natural areas, water bodies and wetlands) and further grouped those classes into anthropized areas (agricultural areas and artificial surfaces) and natural areas (others).

Land cover was included as a random effect in the Generalised Additive Mixed Models to control for differences in landcover types inside and outside PAs.

##### a.3- Elevation

Elevation from the Shuttle Radar Topography Mission (SRTM)^9^ was downloaded via the WorldClim website. Average elevation and slope were extracted for each grid cell.

Elevation was included as a smoothing term in the Generalised Additive Mixed Models to account for biases in elevation of PAs.

##### a.4- Remoteness

We used the map of travel time to cities (in minutes) with data from 2015 at a spatial resolution of approximately one by one kilometre that integrated ten global-scale surfaces that characterize factors affecting human movement rates 8. We extracted the average remoteness of each grid cell.

Remoteness was included as a smoothing term in the Generalised Additive Mixed Models to account for biases in remoteness of PAs.

### Across site analysis –

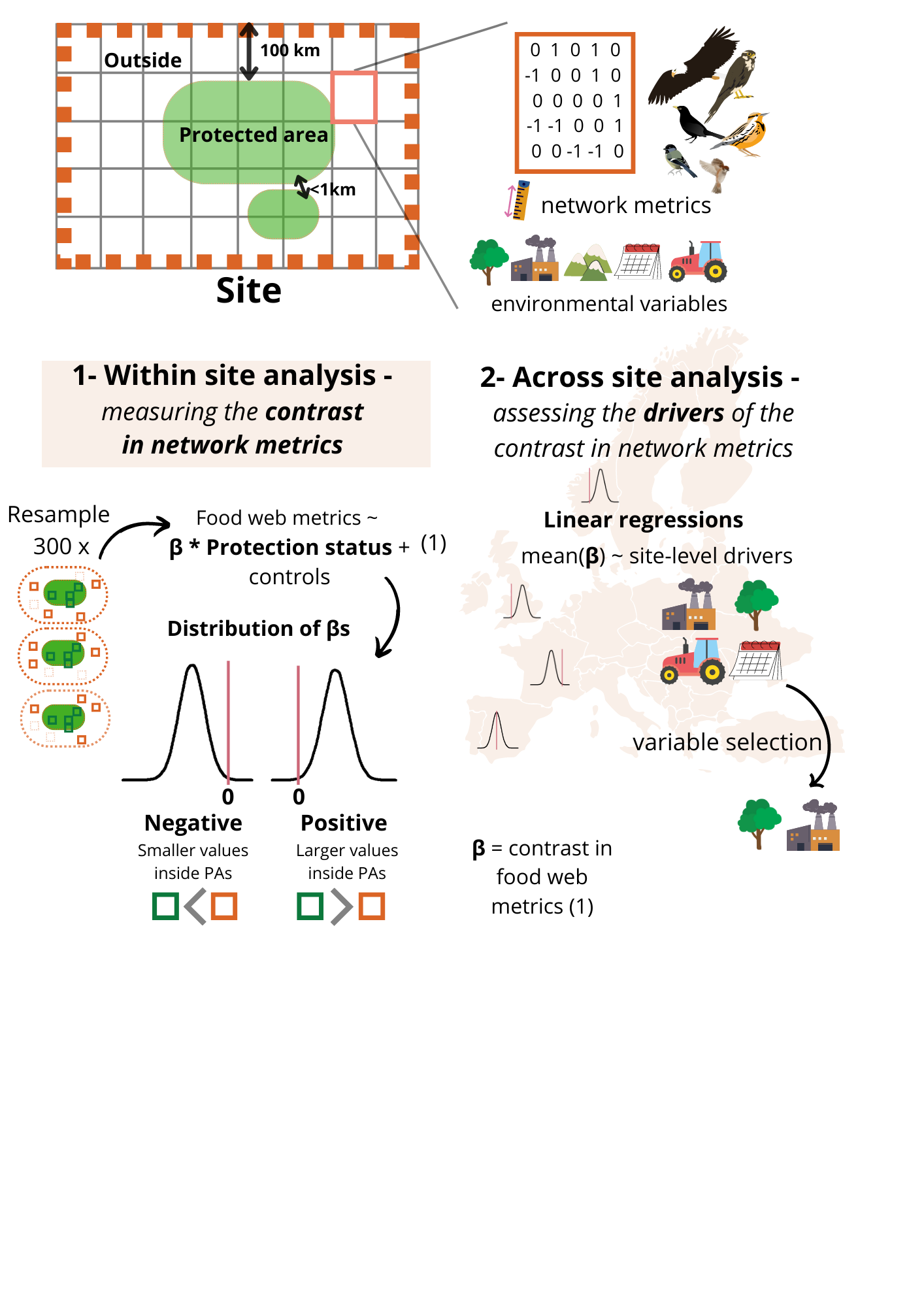

Supplementary Methods Fig. 5: The last step of our analysis was to quantify the role of the environmental context of the sites as an across-site predictor of protection outcome. To quantify this, we ran simple linear models of the mean contrast in food web metrics (mean β) against a suite of environmental predictors, including land cover type, human pressure and protected area characteristics described in Supplementary Methods Table. 3.

To characterise sites in terms of their environmental conditions and types of protected areas, site level characteristics were computed using both environmental already mentioned above and new information such as population density.

Environmental characteristics could be specific to the protected areas, site-wide, or specific to the outside of the protected area. Finally, metrics related to the contrast between the inside and outside were also computed, to have an analogous measure to the contrast in food web metrics, but for environmental conditions. More details about the specific metrics used, their ranges and why they were included in Supplementary Methods Table. 3.

**Supplementary Methods Table 3. Description of environmental characteristics used to describe the conditions inside a site**. Environmental characteristics could be specific to the protected areas, site-wide, or specific to the outside of the protected area. Metrics related to the contrast between the inside and outside were also computed, to have an analogous measure to the contrast in food web metrics, but for environmental conditions. This table describes our biological reasons for including them in the analysis, as well as the observed range across the dataset and interpretation when needed.

| Site-level drivers | Area of calculation | Observed range | Hypothesis | Interpretation |
| --- | --- | --- | --- | --- |
| Average elevation (meters) | Whole site | 34 – 1685 meters | Area of higher elevation are unlikely to be intensively managed and highly anthropized, so might show more pristine environments and thus smaller differences in food web metrics. |  |
| Average slope (degrees) | Whole site | 6e-2 – 0.24 degrees | Area of steeper slope are unlikely to be intensively managed and highly anthropized, so might show more pristine environments and thus smaller differences in food web metrics. |  |
| Average remoteness (minutes) | Whole site | 6 – 388 minutes | More remote sites are expected to show least contrast in food web metrics (inside = outside) as the outside and inside should differ less in terms of disturbance intensity. | Travel time in minutes to the closest city |
| Habitat diversity | Whole site | 0.35 – 1.58 | More habitat diversity could favour higher trophic levels which need more resources to survive at the regional level. | 0 = One single land cover type covering the whole site  5 = Many land cover types in identical proportion |
| Average date of designation (year) | Protected grid cells | 1964 - 2014 | Older protected areas are expected to correlate with a positive contrast in food web metrics (inside > outside), as they have their habitat should have been shielded from disturbance for longer time periods. |  |
| Average protection level | Protected grid cells | 2.4 - 10 | Protected areas with higher protection levels (strict nature reserves, national parcs) are expected to show higher contrast in food web metrics (inside > outside), as they should better shield against disturbances. | 1 = All PAs have lower protection levels  10 = All PAs have higher protection levels |
| Mean protected area | Protected grid cells |  | Larger protected areas should host more species, potentially more specialised species and facilitate movement within protected landscapes, and thus increase contrast in food web metrics (inside > outside) |  |
| Protected area clustering (percentage) | Protected grid cells | 28 – 100% | More fragmented areas are expected to be worse for species richness and specialisation as they will have more edges, probably more fragmented landscapes and might prevent dispersal if the surrounding habitat is unhospitable for certain species ^10^ | 0%= most disaggregated 100% = most aggregated |
| Human density inside protected areas | Protected grid cells |  |  |  |
| Difference in proportion of forested land cover inside the protected areas (percentage) | Protected grid cells |  | Forested protected areas have been shown to benefit forest birds in tropical regions ^1^, thus we expect forested protected areas to show better outcomes in terms of food web metrics too. | 0 – 100% of forested land cover types |
| Proportion of protected areas managed for bird protection | Protected grid cells | 0 – 100% | Protected areas managed for birds are expected to positively increase the contrast in species richness, host less generalist species and longer mean food chain length. | 0 = no protected areas were managed for bird protection  100 = all protected areas were managed for bird protection |
| Proportion of agricultural land cover outside the protected areas (percentage) | Non-protected grid cells | 1 – 96% | Area with more urban area outside protected areas might show higher contrast in food web metrics, as species have less pristine habitat to disperse to ^11^. | 0 – 100% of agricultural land cover types |
| Proportion of forested land cover outside the protected areas (percentage) | Non-protected grid cells | 1 – 96% | Forested areas outside protected areas could benefit species inside protected areas. Thus, we might expect areas with forested areas outside to show either null or positive differences in food web metrics (outside = inside or outside > inside). | 0 – 100% of forested land cover types |
| Proportion of urban land cover outside the protected areas (percentage) | Non-protected grid cells | 1 – 96% | Area with more urban area outside protected areas might show higher contrast in food web metrics, as species have less pristine habitat to disperse to ^11^. | 0 – 100% of urban land cover types |
| Human density outside protected areas | Non-protected grid cells |  | Area with more human density outside protected areas might show higher contrast in food web metrics, as species have less pristine habitat to disperse to ^11^. |  |
| Difference in proportion of urban land cover outside the protected areas (percentage) | Inside - outside |  | The higher the contrast between protected and non-protected grid cells, the higher the expected contrast in food web metrics ^11^. | 0 – 100% of urban land cover types |
| Difference in proportion of agricultural land cover outside the protected areas (percentage) | Inside - outside |  | The higher the contrast between protected and non-protected grid cells, the higher the expected contrast in food web metrics ^11^. For agricultural land cover, we expect a negative relationship between difference in agricultural landcover and contrast in food web metrics, especially for specialised birds. | 0 – 100% of agricultural land cover types |
| Difference in proportion of forested land cover outside the protected areas (percentage) | Inside - outside |  | Areas with more forest cover inside than outside is expected to show a higher contrast in food web metrics (inside > outside). Indeed, a positive difference in forest cover would lead to protected communities being very different from those from its surroundings. Thus, the protected area could host more specialist feeders and also longer mean food chain lengths than their surroundings. | 0 – 100% of forested land cover types |
| Difference in human density | Inside - outside |  | The higher the contrast between protected and non-protected grid cells, the higher the expected contrast in food web metrics ^11^. |  |

8.	Copernicus Land Monitoring Service & European Environment Agency. CORINE Land Cover (CLC). (2018).

9.	Farr, T. G. *et al.* The Shuttle Radar Topography Mission. *Rev. Geophys.* **45**, RG2004 (2007).

10.	Williams, J. C., ReVelle, C. S. & Levin, S. A. Spatial attributes and reserve design models: A review. *Environ Model Assess* **10**, 163–181 (2005).

11.	Cabeza, M. & Moilanen, A. Site-Selection Algorithms and Habitat Loss. *Conservation Biology* **17**, 1402–1413 (2003).
